# Taxonomy-agnostic hyperspectral–morphological phenotyping of fungal pathogen chemical-stress responses using machine learning

**DOI:** 10.1101/2025.11.01.686032

**Authors:** Insuck Baek, Seunghyun Lim, Amelia Lovelace, Sookyung Oh, Masoud Kazem-Rostami, Helen Ngo, Moon S. Kim, Lyndel W. Meinhardt, Lalit Kandpal, Minhyeok Cha, Chansong Hwang, Richard D. Ashby, Ezekiel Ahn

## Abstract

Can standardized chemical stress reveal a reproducible, taxonomy-agnostic stress-response fingerprint that is predictive of sample-source labels (crop-of-isolation) in fungal isolates? Six coffee□associated *Colletotrichum* isolates were profiled using four phenolic□branched compounds and compared them with a previously characterized cacao panel. Quantitative morphology, hyperspectral imaging (HSI), and supervised machine learning (ML) yielded panel-specific fingerprints under uniform, isotropic *in vitro* conditions. Circularity, a measure of edge symmetry, was the most informative morphological feature, and ML classified the crop-of-isolation label (coffee vs cacao, in this panel) with 86.7% accuracy in within-panel cross-validation. HSI detected dose-dependent spectral shifts in a targeted subset of isolates and compounds, including changes near 1930 nm in the short-wave infrared, a moisture-sensitive region that warrants robustness checks (e.g., band masking or preprocessing sensitivity) prior to biochemical attribution. Multi□locus phylogeny showed the coffee isolates are polyphyletic, so the predictive signal should be interpreted conservatively as a taxonomy-agnostic phenotype fingerprint associated with crop background in this mixed-lineage panel, acknowledging that crop labels are partially confounded with phylogenetic structure. We propose a “chemical priors” framework as a working hypothesis, in which long-term environmental exposure may imprint stress-response pathways that become legible under simple, standardized probes. This integrative workflow supports scalable screening of eco-friendly antifungals and sensor-driven decision support for high-throughput phenotype-based screening workflows.

**Highlights:**

- Taxonomy-agnostic workflow integrates hyperspectral imaging and morphology.
- Machine learning predicts crop-source labels with 86.7% accuracy in mixed lineages.
- System exhibits robustness, maintaining >86% accuracy even after feature ablation.
- Non-linear ML captures structure missed by linear stats (1.7% variance).
- Enables rapid, sensor-driven antifungal screening without prior DNA sequencing.

## 1. Introduction

Recent research in high-throughput phenotyping has increasingly demonstrated the utility of non-destructive sensors, particularly hyperspectral imaging (HSI) coupled with machine learning (ML), for evaluating plant-pathogen interactions and early disease detection (Rayhana et al., 2023; Tomaszewski et al., 2023). In particular, HSI and ML have been successfully deployed to detect latent anthracnose infections caused by *Colletotrichum* species in various agricultural commodities, including mango and citrus (Siripatrawan and Makino, 2024; Tang et al., 2023; Velásquez et al., 2024). However, while these technologies are widely used to diagnose infections in planta, their application for quantifying dynamic, compound-induced stress responses in vitro remains largely unexplored. Addressing this gap is critical for managing *Colletotrichum*, which causes severe anthracnose and berry diseases in globally significant perennial crops such as cacao (*Theobroma cacao*) and coffee (*Coffea arabica*). A fundamental challenge is extracting reproducible, predictive signals from observable stress phenotypes, a problem we address here using an integrated spectroscopy and machine learning approach. To this end, the anthracnose pathogen *Colletotrichum* was selected as a model system. While notorious for its broad host range, we selected its interaction with two globally significant perennial crops, cacao and coffee. Their crop-associated isolate panels provide an applied testbed for evaluating whether standardized chemical stress yields predictive phenotype fingerprints under uniform and identical experimental conditions. For decades, the primary defense against such pathogens has been the application of synthetic chemical fungicides. However, this strategy is increasingly challenged by the evolution of fungicide-resistant pathogen strains and growing concerns about the environmental and toxicological effects of chemical residues (Cortaga et al., 2023; Schnabel et al., 2021; Tsalidis, 2022). This creates an urgent need for novel antifungal strategies that are not only effective but also ecologically sustainable and are often sought from natural sources (Cantrell et al., 2012; Thind, 2017). This led us to a central hypothesis: that a pathogen’s observable response to chemical stress can encode a stable, learnable phenotype fingerprint that is useful for screening and classification. To test this, a novel, high-throughput workflow was developed combining quantitative morphology, HSI, ML. We contend that this integrative platform provides a powerful new lens for sensor-driven, non-destructive phenotyping and predictive modeling with practical applications in phytopathology.

The two crop-associated isolate panels provide a practical context to deploy this workflow. Here, we applied our integrative approach to quantify what we call ‘chemical priors’, stress-response phenotypes that are hypothesized to reflect prior exposure histories or persistent physiological states, and can be leveraged as predictive fingerprints. To elicit these phenotypes, we utilized a 2x2 matrix of novel, stabilized phenolic-branched antifungals derived from sustainable feedstocks (Borges et al., 2018; Du et al., 2015; Huang et al., 2021; Kazem-Rostami et al., 2023; Lew et al., 2018). These compounds, with systematic variations in their phenolic headgroups and backbones, allow us to probe the structural basis of antifungal activity, building upon our initial findings in the cacao panel where specific derivatives induced reproducible morphological fingerprints (Ahn et al., 2026).

This observation prompted a central question: can standardized chemical-stress phenotypes support robust, cross-validated prediction of source labels in a mixed-lineage isolate panel? From this question arise two possible explanations. Any predictive signal associated with crop background could reflect host-linked effects, biogeographical history, or lineage structure; therefore, interpretations must remain conservative without species-resolved experimental control. The distinction is critical, as it determines whether a phenotype fingerprint primarily reflects compound-driven stress physiology or panel-specific biological context. Given the differences in host chemistry and cultivation context reported for cacao and coffee systems (Cassamo et al., 2022; de Sousa et al., 2021; Lahive et al., 2019; Nugroho et al., 2016; Winters et al., 2024), it is plausible that fungal isolates sampled from these crops exhibit divergent stress-response phenotypes; however, overlap between agroecosystems and lineage confounding can also generate apparent differences, so we treat crop-of-isolation as an empirical label rather than a mechanistic claim.

Our primary goal was to develop, evaluate, and benchmark an integrated workflow that can quantify and model these signatures. By applying this platform to two crop-associated isolate panels, the specific objectives were to: (1) quantify compound-induced morphological fingerprints; (2) use ML to test their predictive power for crop-of-isolation labels (in this panel); and (3) use HSI to explore the underlying physiological basis of these predictive phenotypes.

## 2. Materials and methods

### 2.1. Biobased compounds

The procedures for the production and characterization of phenolic soybean oil branched fatty acids (PhSOFA) and their ethylenediamine-carrying fatty amide derivatives (PhSOAM) have been previously reported in detail (Huang et al., 2021; Lew et al., 2018). The procedures for the production and characterization of beechwood creosote-branched brown grease fatty acids (BCBGFA) and their ethylenediamine-carrying fatty amide derivatives (BCBGAM) have also been previously reported in detail (Kazem-Rostami et al., 2023). Each of the four final compounds was dissolved in pure ethanol to create stock solutions of 10 mg/mL prior to application.

### 2.2. Fungal isolates and culture conditions

Six coffee-associated fungal isolates of *Colletotrichum* spp. (P23-55, P24-83, P24-85, P24-88, P24-192, and P24-193) were used in this study. The specific origin, sample metadata, and isolation year for each coffee isolate are provided in Table S4. For the comparative analysis, data from a previously characterized set of cacao-associated fungal isolates (including *Pestalotiopsis* sp. CGH5 and *C. gloeosporioides* CGH17 and CGH49) were also included (Ahn et al., 2026). All fungal isolates were routinely maintained on Potato Dextrose Agar (PDA) medium and incubated in the dark at 24°C for 7 days to promote fungal growth.

### 2.3. Phylogenetic analysis

To elucidate the phylogenetic relationships among the fungal isolates, three gene regions were analyzed: the internal transcribed spacer (ITS), glyceraldehyde-3-phosphate dehydrogenase (GAPDH), and actin (ACT). Genomic DNA was extracted from 7-day-old isolates grown on PDA using the Quick-DNA Fungal/Bacterial Miniprep kit (Zymo) according to the manufacturer’s instructions. The ITS, GAPDH, and ACT gene regions were amplified using the primer pairs ITS-1 and ITS-4 (White, 1990), GDF and GDR (Templeton et al., 1992), and ACT-512F and ACT-783R (Carbone and Kohn, 1999), respectively. PCR was performed using the HiFi Phusion polymerase kit (NEB) in 50 µL reactions with 1 µL of genomic DNA as a template. Thermal cycling conditions for ITS amplification consisted of an initial denaturation at 98°C for 3 min, followed by 35 cycles of 98°C for 30 s, 52°C for 30 s, and 72°C for 30 s, with a final extension at 72°C for 10 min. For ACT amplification, an annealing temperature of 61°C and an extension time of 15 s were used. For GAPDH amplification, an annealing temperature of 60°C and an extension time of 15 s were used. PCR products were purified using a QIAQuick PCR kit (Qiagen) and sequenced in both forward and reverse directions at Psomagen (Rockville, MD, USA). Forward and reverse reads for each gene were assembled de novo to generate a consensus sequence for each isolate in Geneious Prime (v2025.2.1), and low-quality bases at the sequence ends were manually trimmed. These newly generated sequences were combined with previously characterized sequences from the cacao-associated isolates and reference sequences from closely related *Colletotrichum* species obtained from the NCBI GenBank database. Phylogenetic analyses were used to contextualize lineage relatedness among isolates; species-level taxonomic assignment was not the primary objective of this study.

Sequences for each gene were aligned using the MUltiple Sequence Comparison by Log-Expectation (MUSCLE) algorithm implemented in Geneious Prime. Phylogenetic inference was conducted for two datasets using MEGA 11: (1) the ITS region alone, and (2) a concatenated dataset of the three gene regions (ITS, GAPDH, and ACT). For each analysis, the best-fit nucleotide substitution model was selected in MEGA 11 based on the lowest Bayesian Information Criterion (BIC) score. For the ITS-only analysis, a phylogenetic tree was constructed using the Neighbor-Joining (NJ) method, with evolutionary distances computed using the Kimura 2-parameter model with a gamma distribution (K2+G). For the concatenated analysis, an NJ tree was also constructed, with evolutionary distances computed using the Tamura-Nei model with a gamma distribution (TN+G). For both trees, nodal support was assessed via bootstrap analysis with 1,000 replications, and bootstrap values of 75% or greater are shown on the corresponding nodes.

### 2.4. Antifungal activity assay

The antifungal effects of the four compounds were tested via a modified agar plate assay. Each compound (10 mg/mL stock) was spread evenly across PDA plates (15 µL, yielding 150 µg/plate) using a sterile spreader. Control plates received 15 µL of pure ethanol. A 5-mm mycelial plug from the growing edge of a 7-day-old culture of each fungal isolate was placed at the center of the treated plates, which were then incubated at 24°C in the dark for 96 hours. Each isolate-treatment combination had at least ten replicates. Each replicate consisted of an independently prepared PDA plate inoculated with a single 5-mm plug (one plate = one biological replicate).

### 2.5. Image acquisition and morphological feature extraction

Following 96 hours of incubation, digital images of the fungal colonies were acquired using a Nikon Z8 camera fitted with a Nikkor Z 20mm f/1.8 S lens (Nikon, Tokyo, Japan), mounted on a Kaiser R2N CP stand (Kaiser Fototechnik, North White Plains, NY). Images were captured under consistent lighting conditions provided by an RB 218N HF lighting unit equipped with two 18-W high-frequency fluorescent lamps (5400 K daylight-balanced, CRI > 90) positioned at approximately 45° incident angles to ensure uniform illumination and minimize specular reflection across the plate surface. Colony morphology was quantified using the SmartGrain software (version 1.1) (Tanabata et al., 2012). The outline of each colony was manually traced within the software by a single trained analyst to ensure consistency. Tracing followed a standardized protocol (fixed magnification and boundary rules) to minimize user-to-user variability, and image filenames were coded to blind the analyst to specific treatment identities during the tracing process. From these outlines, SmartGrain automatically calculated the following morphological traits: colony area (mm²), perimeter (mm), length (mm), width (mm), LWR, circularity, and the asymmetry index defined as the distance between the intersection of length and width and the center of gravity (IS & CG, in mm).

### 2.6. Statistical analysis and machine learning evaluation

General statistical analyses were performed using JMP Pro 17 (SAS Institute Inc., Cary, NC, USA) (Klimberg, 2023). To assess differences in morphological traits between treatment groups, a one-way Analysis of Variance (ANOVA) was performed, followed by a Tukey’s Honestly Significant Difference (HSD) post-hoc test for all pairwise comparisons. For multivariate analyses, all quantitative morphological measurements were log-transformed to stabilize variance and better approximate a normal distribution. PCA was performed on the standardized, log-transformed data. Pearson’s correlation analysis was used to assess linear relationships between pairs of the transformed traits. PCA and PERMANOVA were first conducted on the coffee-only dataset to evaluate chemical and isolate effects (Fig. 3), and then repeated on the combined coffee + cacao dataset to evaluate crop-of-isolation label and treatment effects (Fig. 5). Differences in multivariate morphological profiles between groups were tested using PERMANOVA based on Euclidean distances of the log-transformed traits, with 9,999 permutations. Additionally, to dissect the interaction between isolate identity and chemical treatment, a two-way factorial ANOVA was performed on log-transformed traits.

For the comparative analysis, the coffee isolate dataset was merged with the previously published cacao-associated isolate dataset (Ahn et al., 2026), which was generated using the identical experimental setup, media preparation protocol, and imaging equipment to minimize batch effects. To predict whether a sample’s crop-of-isolation label (cacao vs coffee, in this dataset) could be predicted from its morphological response, a comprehensive suite of classification models was evaluated using the “Model Screening” platform in JMP Pro 17. Models were trained using JMP Pro 17 “Model Screening” with default settings for benchmarking; detailed model settings are provided in Supplementary Data 1 (model reports). Due to an unequal number of samples between the two panels, the dataset was balanced by randomly subsampling the larger class to contain 180 samples from each crop-of-isolation, using a fixed random seed (seed = 1), an approach chosen to preserve the original data distribution and avoid introducing artifacts from oversampling techniques. The predictor variables included treatment type and the seven morphological traits, which were standardized prior to model fitting. Pathogen isolate identity was deliberately excluded as a predictor to ensure the model learned to classify based on the interaction phenotype rather than memorizing individual isolates. Model performance was assessed using a repeated K-Fold cross-validation strategy (*k* = 5, random seed = 1). Because multiple samples per isolate were present, cross-validation performance is primarily interpreted as within-panel predictability. To assess robustness to isolate-level confounding, we additionally performed isolate-grouped cross-validation using StratifiedGroupKFold with strain as the grouping variable (*k* = 2 after excluding CGH5, due to the limited number of cacao strains), as reported in Supplementary Data 1. Primary analysis was performed in JMP Pro 17; additional sensitivity analyses (grouped CV) were performed using Python scripts, with outputs provided in Supplementary Data 1. To address potential taxonomic and class-composition confounding, we conducted two additional sensitivity analyses: (1) excluding the highly responsive cacao isolate (CGH5) to assess whether classification relied solely on extreme phenotypes, and (2) removing the top predictor (circularity) to test feature redundancy.

### 2.7. Hyperspectral imaging and spectral analysis

To investigate the early physiological responses induced by antifungal treatment, HSI was conducted on 96-hour incubated colonies of *Colletotrichum* spp. Fungal cultures were initially grown on potato dextrose agar (PDA) plates for 7 days at 24 °C in the dark. From these mature cultures, 5 mm diameter plugs were collected using a sterile biopsy punch and transferred to fresh PDA plates containing one of two phenolic-branched compounds, PhSOAM or BCBGAM, selected as a representative subset based on their superior antifungal potency in the initial screening, applied at three concentrations (150, 300, and 450 µg per plate). Control plates were treated with ethanol only. Three fungal isolates were selected for HSI analysis based on their distinct morphological responses observed in prior antifungal assays. The cacao-associated isolate CGH5 (*Pestalotiopsis* sp.), which previously exhibited dramatic reductions in colony area upon PhSOAM treatment, was included as a representative of a highly responsive genotype. Two coffee-associated *Colletotrichum* isolates, P24-85 and P24-192, were selected to capture intra-panel variation in chemical response. For each isolate and treatment condition, four biological replicates were prepared, with each replicate consisting of a single, independently cultured fungal plug. HSI was carried out using custom-built line-scan systems as previously described (Kim et al., 2011). In this study, three kinds of imaging modalities were employed: fluorescence imaging, VNIR reflectance imaging, and SWIR reflectance imaging. For VNIR reflectance (401.5–1001 nm spectral range) and fluorescence imaging (438–718 nm spectral range), the system was equipped with an electron-multiplying charge-coupled device (EMCCD) camera (Luca DL 604M, Andor Technology, South Windsor, CT, USA) coupled to an imaging spectrograph (Headwall Photonics Inc., Fitchburg, MA, USA). Reflectance illumination was provided by four 150-W quartz tungsten-halogen lamps powered by a DC supply, while fluorescence excitation was achieved using two UV-A LED light sources centered at 365 nm. For SWIR reflectance imaging (842–2532 nm spectral range), a mercury cadmium telluride (MCT) array detector coupled with a SWIR spectrograph (Micro-Hyperspec SWIR, Headwall Photonics, Fitchburg, MA, USA) was used, illuminated by gold-coated halogen lamps to maximize infrared output and minimize thermal drift. All plates were scanned in their entirety, and spectral data were extracted from consistent, standardized regions of interest (ROIs) within each fungal colony. Spectral signatures were calibrated using white reference and dark signal, and F-value analysis was applied to identify wavelength regions showing statistically significant variation as a function of treatment and strain identity through one-way ANOVA F-statistics. Further details on system specifications and spectral calibration procedures are provided in Baek et al. (2025). Because the SWIR region near ∼1930 nm is moisture-sensitive, interpretation focuses on treatment-associated spectral variation rather than biochemical assignment without explicit water-effect modeling.

## 3 Results

### 3.1 Potent but distinct effects of amide derivatives PhSOAM and BCBGAM

The two ethylenediamine-carrying amide derivatives, PhSOAM and BCBGAM, were among the most potent growth inhibitors tested. PhSOAM exhibited potent but selective antifungal activity, reducing colony area in the most susceptible isolates, P24-192 and P24-85, by 19.4% (*p* < 0.001) and 20.7% (*p* < 0.001), respectively (Fig. S1A). This reduction is visually illustrated for P24-192 (Fig. S1B, C). In contrast, BCBGAM induced slightly stronger inhibition in certain isolates (e.g., 20.1% in P24-192) but was less effective against others (Fig. 1A). While both amides reduced size metrics, their shape metrics (circularity, LWR, IS&CG) diverged, underscoring a size–shape decoupling that motivates the downstream multivariate and predictive analyses. PhSOAM triggered diverse morphological responses, in some cases inducing irregular expansion (e.g., a decrease in circularity in isolate P24-193) (Fig. S1A). Conversely, BCBGAM promoted a more compact, rounded morphology in three of the six isolates (Fig. 1A). The most striking response was observed in isolate P24-192, where BCBGAM treatment induced a significant increase in colony elongation (LWR), as illustrated in the representative images (Fig. 1).

**Fig. 1.**
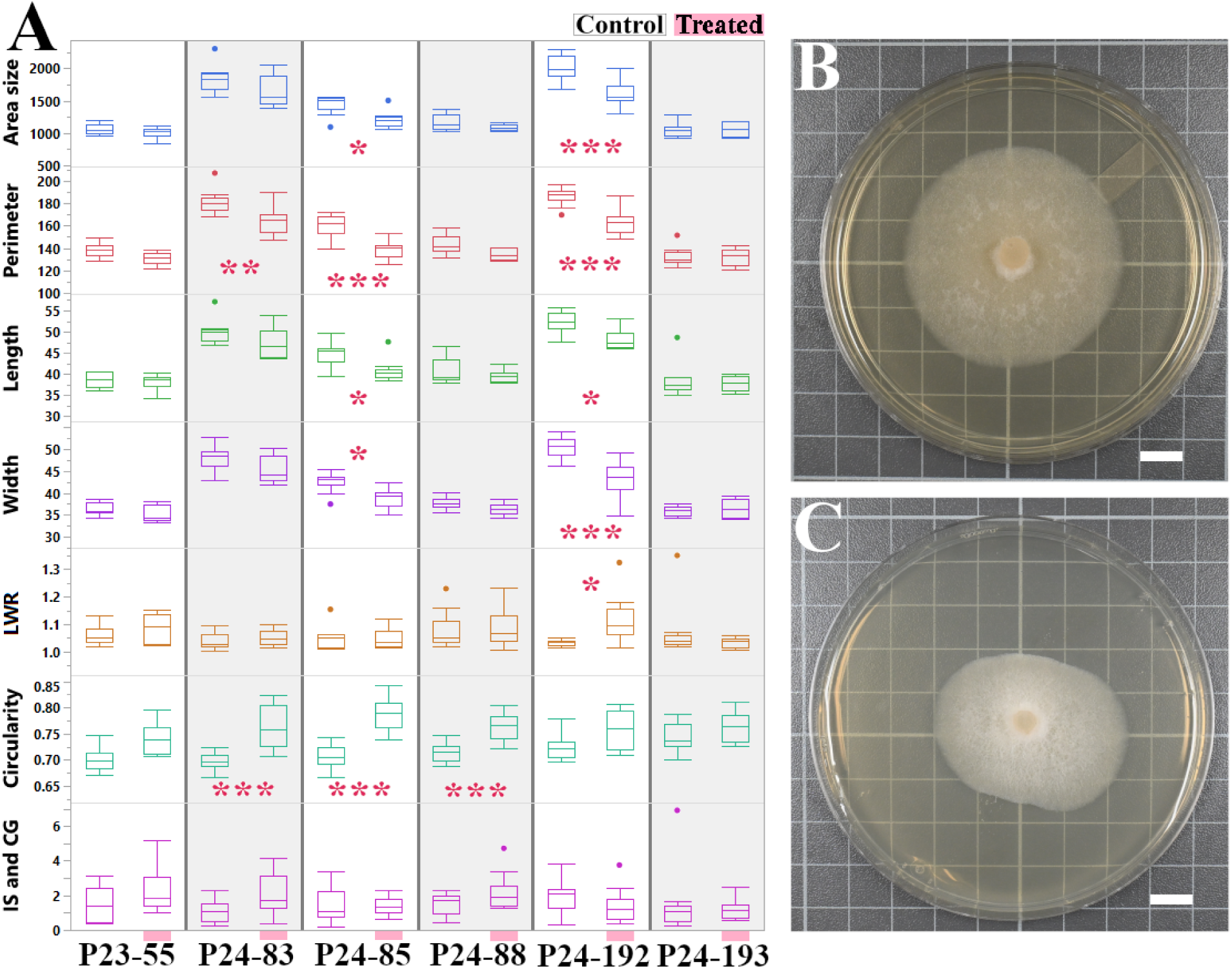
Representative morphological disruption by phenolic antifungal compounds, illustrated using BCBGAM-treated isolates, including P24-192. (A) Box plots quantifying the morphological response of six *Colletotrichum* sp. isolates to one representative phenolic antifungal (BCBGAM), which exhibited isolate-specific effects. BCBGAM is featured because it elicited the most visually distinct morphological changes in isolate P24-192. While PhSOAM showed stronger overall inhibition (Fig. S1), the response to BCBGAM better illustrates the complex ‘morphological fingerprint’ central to this study, highlighting our focus on quantifying reproducible stress-response fingerprints for downstream screening and predictive modeling, rather than solely quantifying growth inhibition. Measured traits include colony area (mm²), perimeter (mm), length (mm), width (mm), LWR, circularity, and an asymmetry distance (IS & CG, mm). Significant differences between a treated group and its corresponding control (Tukey’s HSD; \**p* < 0.05; \*\**p* < 0.01; \*\*\**p* < 0.001) are marked with asterisks. (B-C) Representative images of isolate P24-192 after 96 hours on (B) a control plate and (C) a plate treated with BCBGAM. Sample size, n = 10 for all groups except P24-192 and P24-193 controls (n = 11). Scale bars = 1 cm.

### 3.2 Variable and opposing effects of fatty acid analogs

The two phenol-branched fatty acids, PhSOFA and BCBGFA, exhibited overall weaker antifungal activity than their corresponding amide derivatives, yet each triggered highly distinctive and isolate-specific morphological responses (Fig. 2; Fig. S2–S3).

**Fig. 2.**
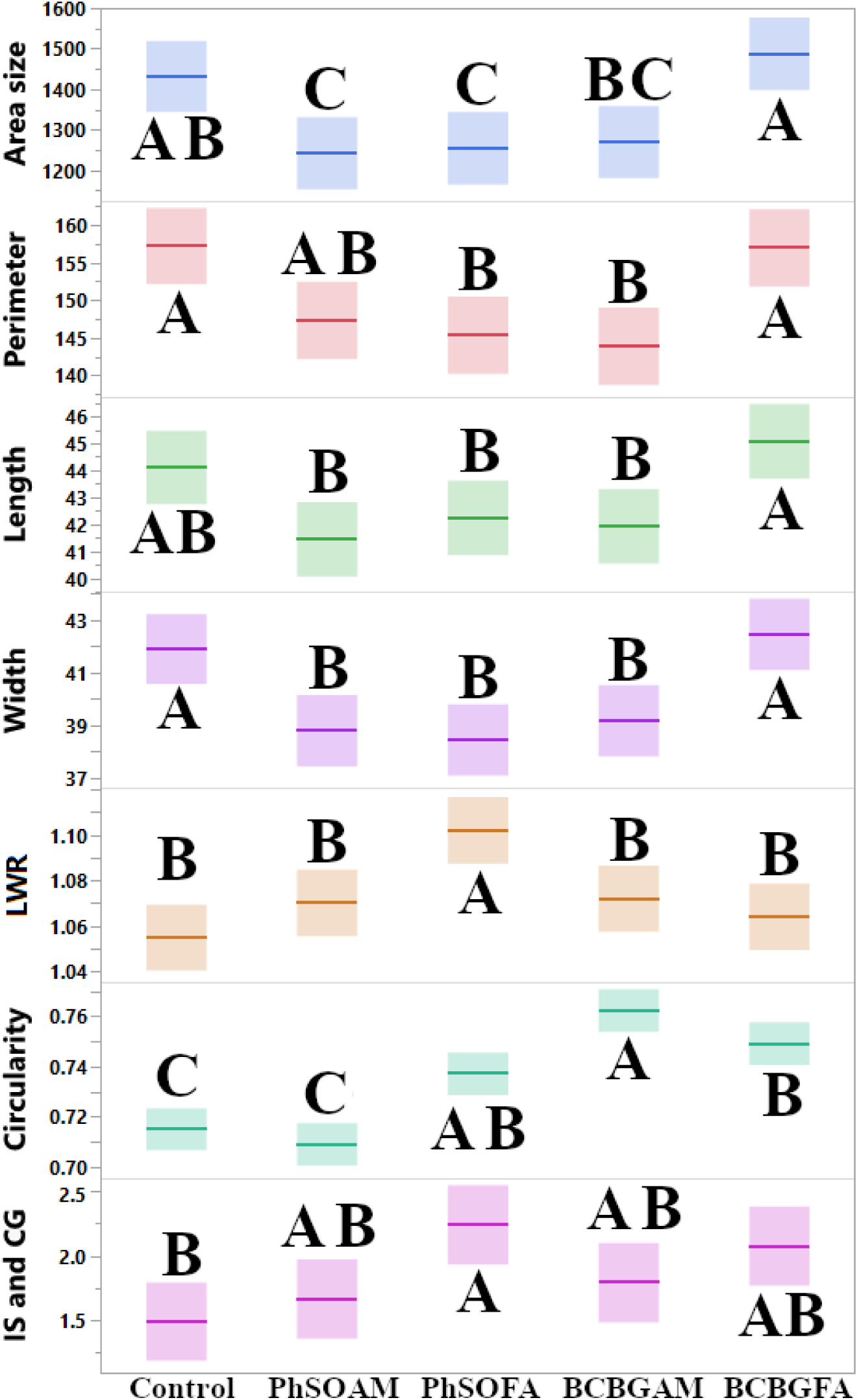
Comparative analysis of the antifungal effects of four phenolic-branched compounds on coffee-associated *Colletotrichum* isolates. Box plots show the pooled morphological data from six *Colletotrichum* isolates under four treatment conditions (PhSOAM, BCBGAM, PhSOFA, and BCBGFA) and an untreated control. Measured traits include colony area (mm²), perimeter (mm), length (mm), width (mm), LWR, circularity, and an asymmetry distance (IS & CG, mm). Different letters denote statistically significant groupings (*p* < 0.05) determined by Tukey’s HSD.

PhSOFA demonstrated selective growth inhibition, significantly reducing colony size in two of the six isolates: P24-83 and P24-192 (Fig. S2A). For example, PhSOFA treatment reduced the colony area of isolate P24-83 by 18.8% (*p* < 0.01), a change illustrated in the representative images (Fig. S2B–C). In contrast, BCBGFA’s effects were highly variable. It had no significant inhibitory effect on any isolate but paradoxically promoted the growth (assessed by area size measurement) of P24-85 by 16.5% (*p* < 0.05), as detailed in Fig. S3. This growth stimulation was consistent across replicates, suggesting a potential hormetic response or isolate-specific metabolic capability to metabolize the fatty acid substrate, rather than an experimental artifact. The remaining isolates were largely resistant to either compound. Morphological shape metrics highlighted further, though more sporadic, divergence between the two treatments. The effects on shape metrics were more sporadic: significant increases in colony circularity were rare, observed only for BCBGFA in P24-83 and PhSOFA in P24-193, while PhSOFA was the only compound to cause significant increases in both colony elongation (LWR) and asymmetry (IS & CG), effects that were limited to isolate P24-192 (Figs. S2 & S3).

When morphological data were pooled across all isolates (Fig. 2), a clear hierarchy of antifungal potency within this plate assay, based on the reduction in colony area, was observed. PhSOAM remained the most potent growth inhibitor. It was followed by PhSOFA. The other amide, BCBGAM, showed no significant difference from the control, while BCBGFA was the least effective, showing no significant inhibition. Despite variable effects on colony size, the compounds induced reproducible shifts in shape. PhSOFA uniquely elevated the LWR. A significant increase in colony circularity was observed for three compounds: PhSOFA, BCBGAM, and BCBGFA. Furthermore, only PhSOFA caused a significant increase in the asymmetry distance (IS & CG), pointing to non-lethal but geometry-altering cellular effects. Together, these data show isolate- and structure-dependent effects across compounds (Fig. 2; Figs. S2–S3). All isolate × compound × trait comparisons were evaluated and reported as *p*-values in Supplementary Data 1, enabling exhaustive auditing of treatment effects. Two-way ANOVA confirmed significant main effects of strain and chemical treatment and a significant strain × chemical interaction for both log(area) and circularity (Supplementary Data 1).

### 3.3 Multivariate analysis reveals a dominant treatment effect on fungal morphology

PCA of the morphological traits showed that the first two PCs captured 76.2% of the total variance (Fig. S4). However, both chemical treatments and fungal isolates showed considerable overlap in PCA space, indicating that linear ordination provided limited separation between groups. This necessitated the use of supervised machine learning to test for nonlinear, diagnostic patterns. Permutational multivariate analysis of variance (PERMANOVA) within the coffee dataset indicated a small but significant treatment effect and a stronger isolate effect (Compound: pseudo-F = 4.06, R² = 0.0518, *p* = 0.001; Strain: pseudo-F = 13.00, R² = 0.180, *p* < 0.0001). This partial grouping in PCA, combined with substantial overlap, suggests that linear ordination provides limited separation. Because PCA did not fully resolve isolates or crop-of-isolation labels in the combined dataset, we next used supervised ML to test for nonlinear, diagnostic patterns.

Further analyses were conducted to explore the relationships between the seven morphological traits (Fig. S5). The PCA loading plot (Fig. S5A) and partial contributions plot (Fig. S5B) confirmed that the primary axis of variation, PC1, was driven by the four main size metrics (area, perimeter, length, and width). Shape-related traits had greater impacts on subsequent components: LWR and IS & CG contributed most to PC2, while circularity was a major driver of PC3. This clear distinction between size and shape was corroborated by hierarchical clustering, which grouped the four size metrics into one tight cluster and the three shape metrics into another (Fig. S5C). Pearson’s correlation analysis further quantified these relationships, revealing strong positive correlations among all size-related traits (*r* > 0.9 for most pairs, detailed in Supplementary Data 1).

### 3.4 Phylogenetic analysis reveals coffee-associated isolates are polyphyletic and intermixed with cacao-associated isolates

To determine the evolutionary relationships among the isolates, we first conducted a multi-locus phylogenetic analysis using a concatenated dataset of the internal transcribed spacer (ITS), glyceraldehyde-3-phosphate dehydrogenase (GAPDH), and actin (ACT) gene regions (Fig. 3). The analysis revealed the six coffee-associated isolates were polyphyletic, distributing across several distinct clades rather than forming a monophyletic group.

Specifically, isolate P24-88 formed a sister group with *C. karstii* (79% bootstrap support), and this clade in turn was a sister group to isolate P24-193. Isolate P23-55 was placed in a separate, distant clade with *C. pseudoboninense*. The remaining three coffee isolates were located in different parts of the tree: P24-192 clustered with *C. siamense* (99% bootstrap), P24-83 was a sister taxon to *C. rhizophorae* and *C. fructicola*, and P24-85 was placed near *C. gloeosporioides* and *C. kahawae*. This polyphyletic distribution indicates that the coffee isolates used in this study represent multiple distinct lineages within the *Colletotrichum* genus.

An analysis based on the standard fungal barcode, the ITS region, was also performed to directly compare the coffee isolates with cacao-associated isolates (Fig. S6). This analysis further highlighted the complex relationships between isolates from the two panels. Notably, a single, strongly supported clade (99% bootstrap) contained an admixture of isolates from different crop-of-isolation labels. This inclusive clade grouped coffee-associated P24-192, P24-85, and P24-83 together with cacao isolates (CGH17, CGH53) and shade tree isolates (CGH34, CGH38). Overall, based on the ITS region, some coffee isolates are more closely related to cacao-associated isolates than to other coffee isolates, emphasizing mixed lineage structure within the panel, such as P24-88 and P24-193, which formed a separate, well-supported clade (96% bootstrap).

### 3.5 Comparative multivariate analysis of coffee and cacao panels

To investigate whether the chemical compounds induced panel-associated morphological responses, the coffee isolate dataset was combined with a previously generated dataset from cacao-associated isolates [16], and a comparative PCA was performed on the combined, log-transformed data.

The first two principal components accounted for 78.6% of the total variance (PC1: 56.8%, PC2: 21.8%) (Fig. 4). When the data were colored by the sample crop-of-isolation label (Fig. 4A), no clear separation was evident. The coffee (black) and cacao (red) samples were heavily intermixed. Crucially, PERMANOVA revealed that the crop-of-isolation label explained only ∼1.7% of the total variance (pseudo-F = 8.49, R² = 0.0174, *p* = 0.0017). This confirms that linear multivariate methods are insufficient to resolve these phenotypes, justifying the use of non-linear machine learning.

**Fig. 3.**
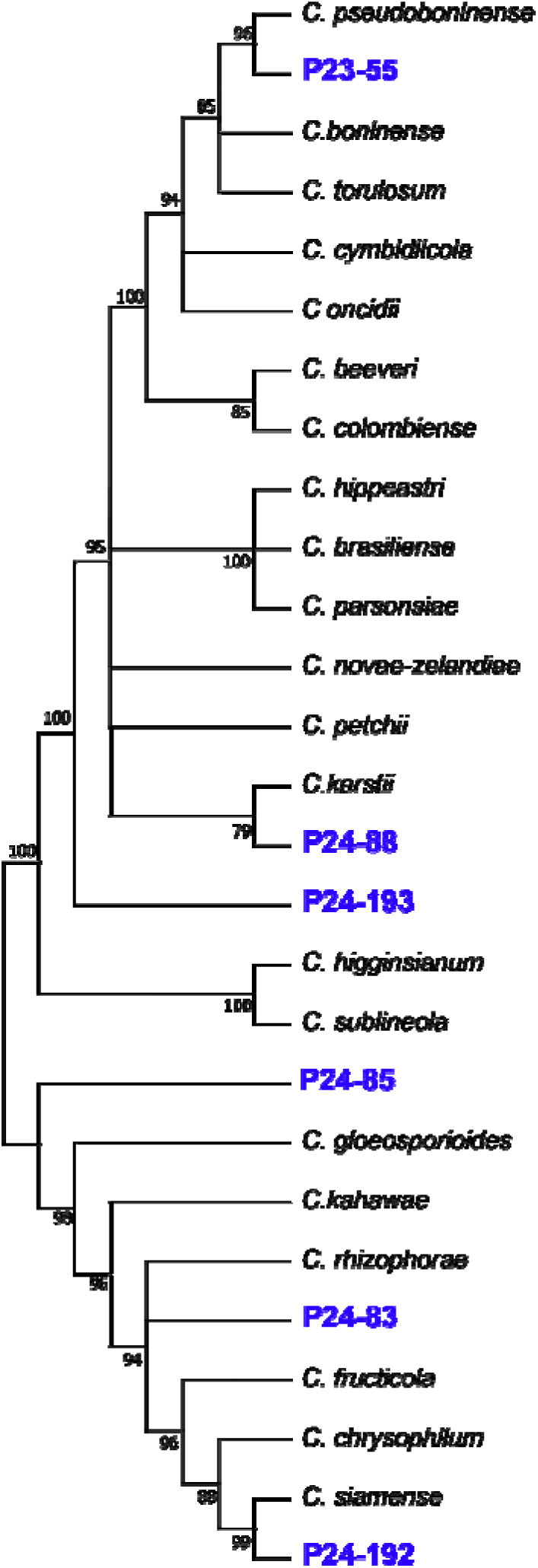
Multi-locus phylogeny reveals the polyphyletic nature of coffee-associated *Colletotrichum* isolates. The phylogenetic tree was inferred by the Neighbor-Joining method using a concatenated alignment of the ITS, GAPDH, and ACT sequences. The evolutionary distances were computed using the Tamura-Nei (TN+G) model. Numbers at the nodes represent bootstrap support values based on 1,000 replications; only values of 75% or greater are shown. The six coffee-associated isolates (highlighted in blue) are shown to be polyphyletic, distributing across multiple distinct clades and indicating affiliations with different known *Colletotrichum* species.

**Fig. 4.**
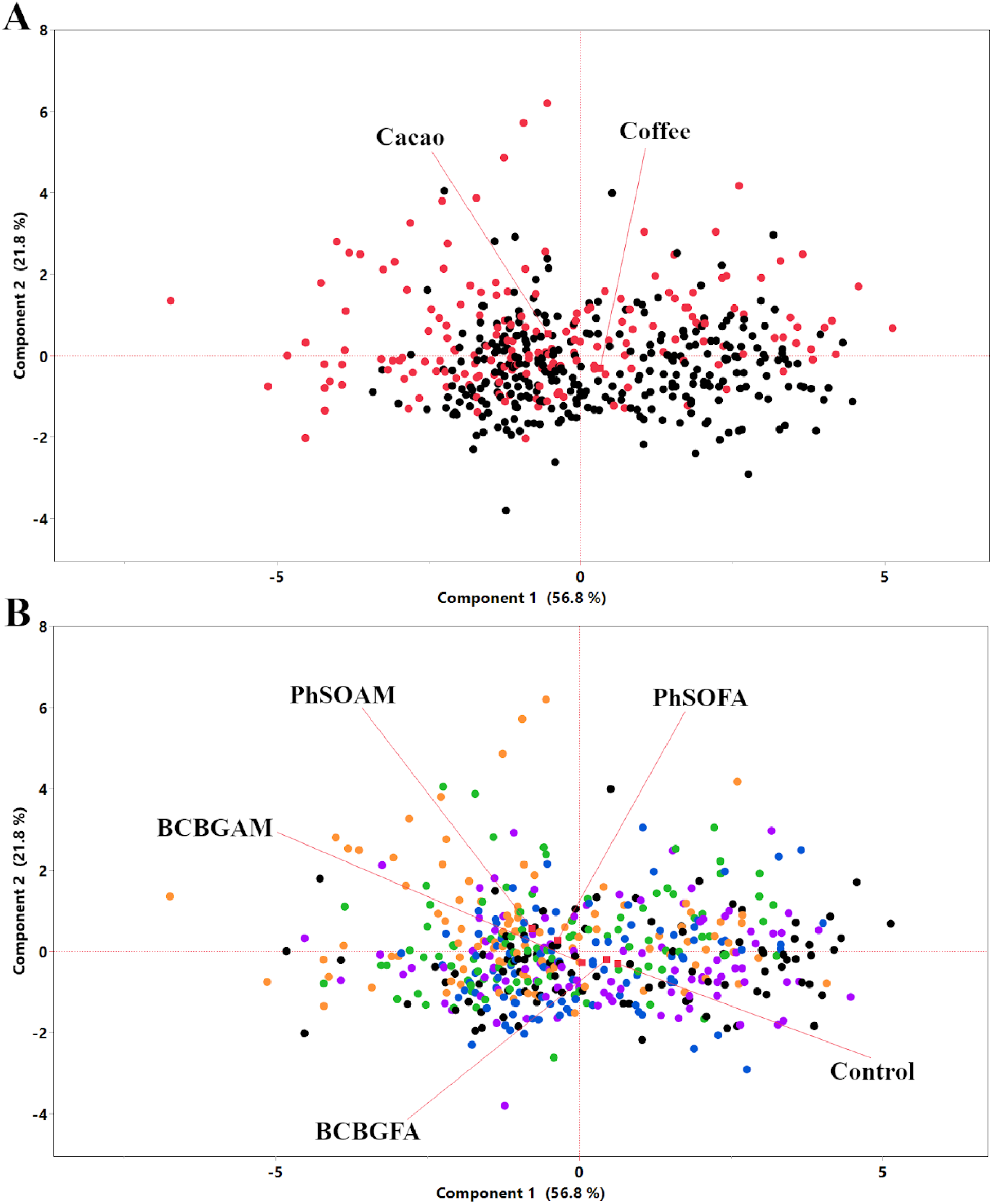
Comparative PCA of coffee and cacao panels reveals treatment-driven separation but panel overlap. PCA of morphological traits from the combined coffee and cacao isolate datasets (log-transformed). (A) PCA scores plot with samples color-coded by panel: Cacao isolates (red) and Coffee isolates (black). Note the significant overlap between the two panels, indicating the first two principal components cannot separate them. (B) The identical PCA scores plot with samples color-coded by chemical treatment: Control (black), PhSOAM (orange), PhSOFA (green), BCBGAM (blue), and BCBGFA (purple). The red squares indicate the mean of each panel in (A) and each chemical treatment group in (B). PERMANOVA indicated a weak crop-of-isolation label effect (Crop-of-isolation: pseudo-F = 8.49, R² = 0.0174, *p* = 0.0017) and a small but significant treatment effect (Compound: pseudo-F = 5.57, R² = 0.0447, *p* < 0.001), both explaining only a small proportion of total variance.

When visualized by treatment type, the analysis showed significant overlaps between the groups (Fig. 4B). The untreated control samples clustered closely with those treated with BCBGFA. A separate cluster was formed by samples treated with PhSOAM and PhSOFA, while the BCBGAM-treated samples were positioned between these two main groupings. This strong overlap indicates that the morphological signatures distinguishing the two panels, if they exist, are not linearly separable and thus are too subtle for PCA to resolve.

### 3.6 Machine learning models classify crop-of-isolation labels from chemical-stress morphology

Following the PCA, which could not clearly separate the two panels, a suite of ML models was employed to quantitatively test if a predictive morphological signature of crop-of-isolation labels (coffee vs cacao, in this dataset) existed within the combined dataset. The analysis revealed that several models could classify crop-of-isolation labels with high accuracy, supporting the existence of a learnable phenotype fingerprint associated with the crop-of-isolation label in this mixed-lineage panel based on the response to the chemical compounds (Fig. 5). A comprehensive screening of nine different algorithms identified the Neural Boosted model (an F1-score of 0.866, and a Matthews Correlation Coefficient (MCC) of 0.733) as the top performer, achieving a validation accuracy of 86.7%, which was followed closely by Nominal Logistic and Generalized Regression (Lasso) models (both ≈85%), as well as Support Vector Machines, Bootstrap Forest, and Fit Stepwise (all 84%) (Table S1).

**Fig. 5.**
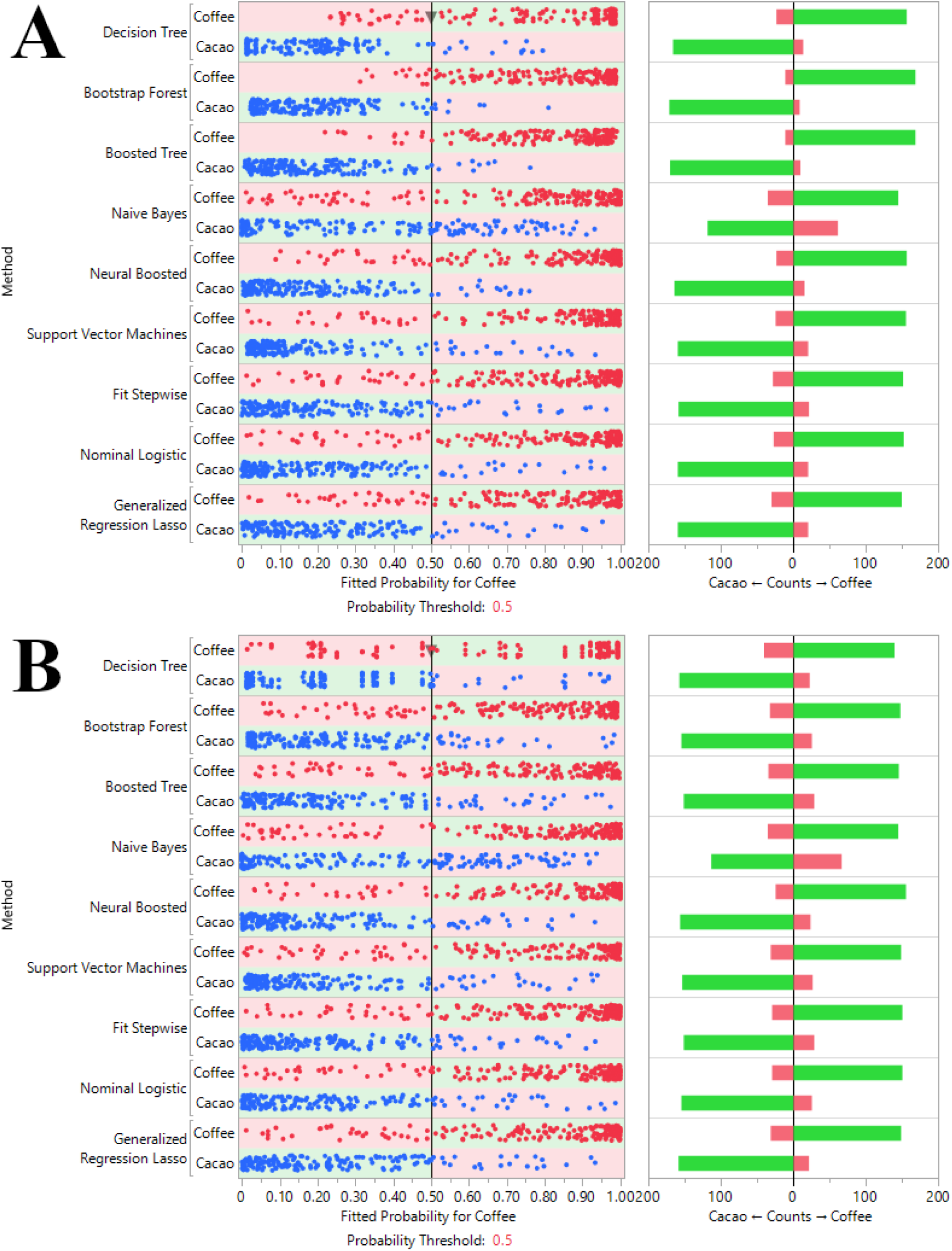
Machine learning classification of crop-of-isolation label based on chemical-induced morphology. Decision thresholds plot showing the performance of nine ML models in classifying a pathogen’s crop-of-isolation labels (cacao vs. coffee) based on its morphological fingerprint from chemical-induced morphology (A: Training set, B: Validation set). Blue dots represent isolates from the cacao panel, and red dots represent isolates from the coffee panel. The plot shows the aggregated results from the k-fold cross-validation, with the confusion matrix on the right summarizing classification performance at a probability threshold of 0.5. ‘Neural□Boosted’ denotes the JMP Pro□17 boosted neural network; default base learner with boosting enabled; repeated 5□fold CV with fixed random seed. Performance is reported for repeated 5-fold cross-validation on a class-balanced dataset; generalization to unseen isolates was not directly tested here.

To understand which traits were driving this successful classification, feature importance was analyzed using the top-performing Neural Boosted model (Table S2). The analysis revealed that circularity was the single most dominant predictor of the crop-of-isolation label, with a Total Effect score of 0.592, which was followed by the primary size-related traits of area (Total Effect = 0.405) and perimeter (Total Effect = 0.168). This indicates that the model primarily learned to distinguish the two crop-associated panels by identifying a unique ‘shape signature’, driven by how circular a colony becomes in response to chemical stress, complemented by information about the colony’s overall size.

To further test the robustness of the classification, the analysis was repeated on the log-transformed dataset after removing circularity, the top predictor (full details in Supplementary Data 1). Even without this primary feature, the models maintained high predictive power. The Neural Boosted model was again the top performer with a validation accuracy of 86.1% (a negligible drop from 86.7% in the full model), followed by the Nominal Logistic (85%), Lasso (85%), and Fit Stepwise (84%) models. Feature importance analysis for this new model revealed a clear shift in predictive strategy; in the absence of circularity, the model pivoted to a ‘size signature’, with area size (Total Effect = 0.825) and perimeter (Total Effect = 0.467) emerging as the most dominant predictors (Supplementary Data 1). Note that because feature importance scores are normalized to sum to 1, the exclusion of the dominant shape feature (circularity) mathematically necessitates a rescaling of the remaining features, resulting in increased weights for size metrics.

Furthermore, a sensitivity analysis excluding the distinct outlier isolate (CGH5) yielded a validation accuracy of 83.5% (Supplementary Data 1), confirming that the predictive signal is not driven solely by a single highly responsive genotype. Finally, to rigorously assess potential data leakage, we performed an isolate-grouped cross-validation using StratifiedGroupKFold with strain as the grouping variable (*k* = 2 after excluding CGH5). Under this stricter evaluation, mean accuracy ranged from 79.3% to 82.9% with MCC values of 0.59–0.66 across the evaluated classifiers (Supplementary Data 1). This grouped evaluation included 6 coffee strains and 2 cacao strains (CGH17 and CGH49), with folds defined at the strain level, confirming that a detectable panel-level phenotype signal remains even when controlling for isolate-level data leakage.

### 3.7 Dose-dependent hyperspectral fingerprints in CGH5 under PhSOAM treatment

To probe the spectral correlates of these morphological fingerprints, we first employed three modalities of HSI: fluorescence HSI, visible and near-infrared (VNIR) reflectance imaging, and short-wave infrared (SWIR) reflectance imaging. We began by testing the response of *Pestalotiopsis* sp. CGH5, a cacao-derived isolate known to exhibit a dramatic 48.8% reduction in fungal growth upon PhSOAM treatment [16]. HSI revealed distinct, dose-dependent responses of *Pestalotiopsis* sp. CGH5 to PhSOAM treatment across all three spectral modalities (Fig. 6).

**Fig. 6.**
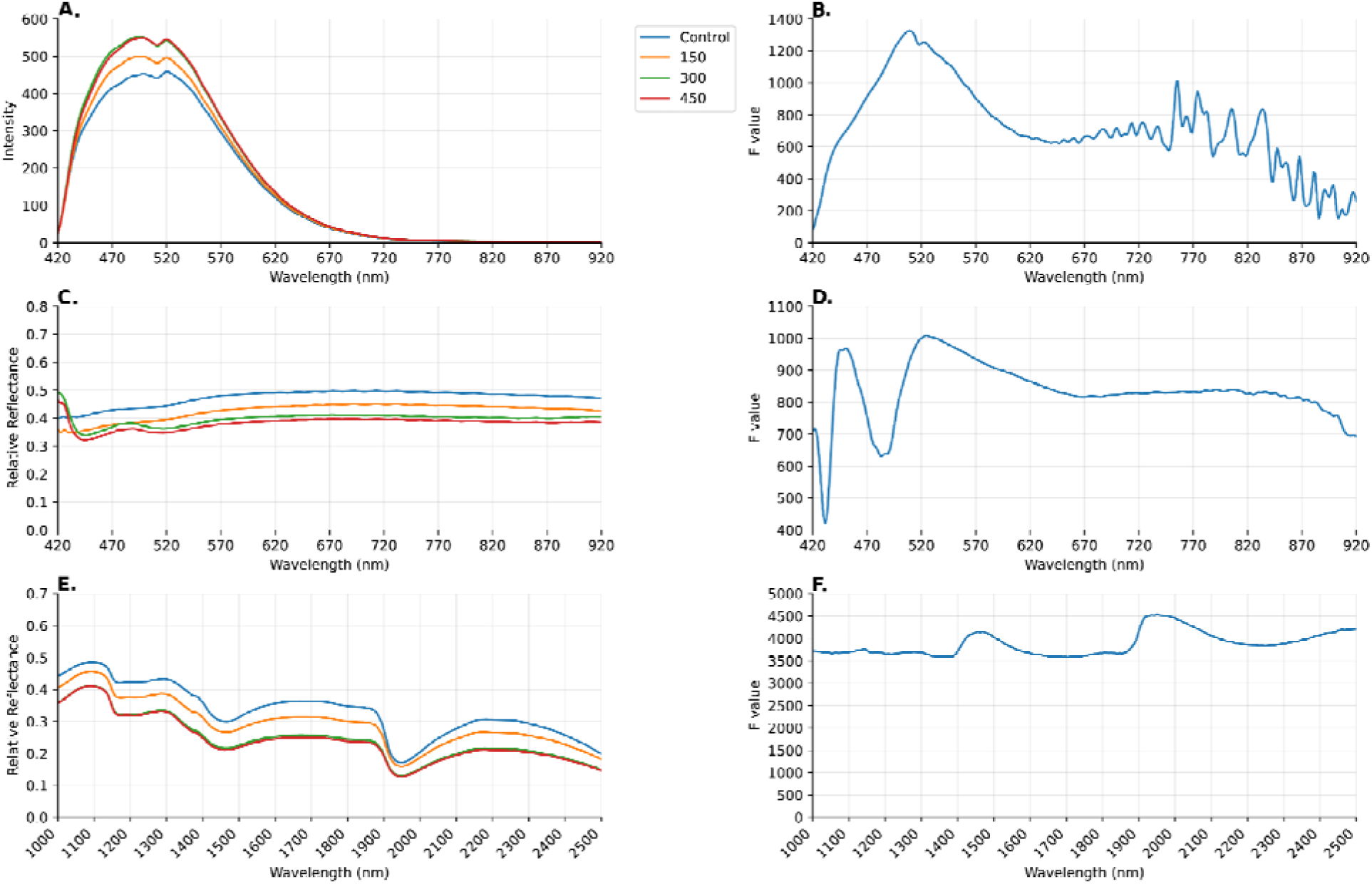
Hyperspectral imaging of *Pestalotiopsis* sp. CGH5 treated with the biobased fungicide PhSOAM across three spectral modalities. (A) Fluorescence intensity profiles of CGH5 colonies treated with 0, 150, 300, and 450 µg of PhSOAM; color indicates dose (control=blue, 150 µg = orange, 300 µg = green, 450 µg= red). (B) F-value curve corresponding to the fluorescence data. (C) Reflectance intensity curves under broadband illumination. (D) F-value profile corresponding to the reflectance data. (E) SWIR reflectance curves showing spectral shifts. (F) F-value curve corresponding to the SWIR data. All data represent the mean of four biological replicates per treatment (n = 4).

### 3.8 Attenuated hyperspectral response in a tolerant coffee isolate

In contrast to the strong response of CGH5, the more tolerant coffee isolate P24-85 exhibited an attenuated and nonlinear hyperspectral response to PhSOAM (Fig. 7). This included a non-monotonic fluorescence signal and muted shifts in the SWIR moisture-sensitive region near ∼1930 nm. These attenuated spectral changes indicate reduced treatment-associated spectral variation in this isolate under the same imaging conditions.

**Fig. 7.**
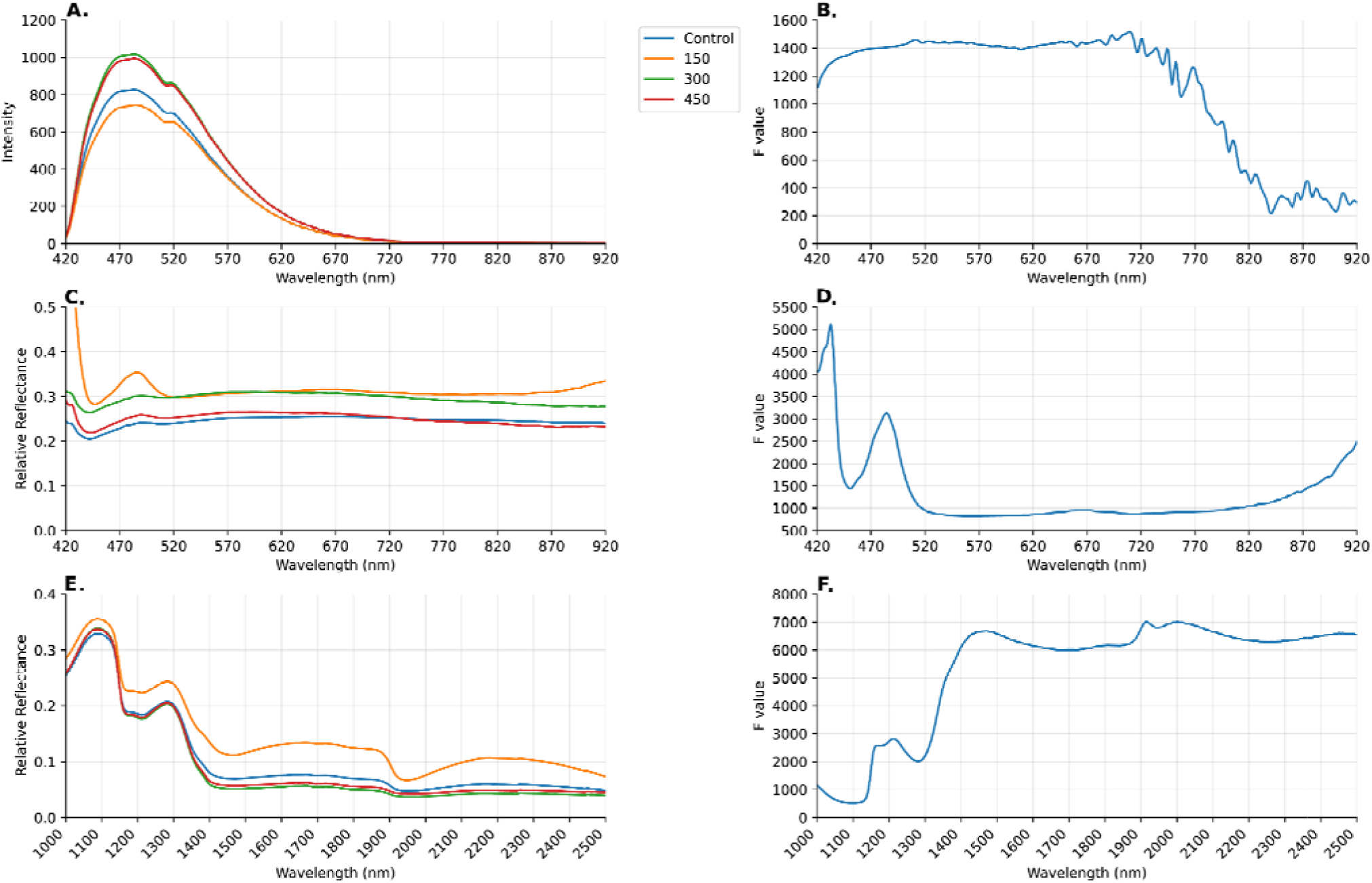
Hyperspectral imaging of *Colletotrichum* sp. P24-85 treated with the biobased fungicide PhSOAM across three spectral modalities. (A) Fluorescence intensity profiles of P24-85 colonies treated with 0 µg (control, blue), 150 µg (orange), 300 µg (green), and 450 µg (red) of PhSOAM. (B) F-value curve corresponding to the fluorescence data. (C) Reflectance intensity curves under broadband illumination. (D) F-value profile corresponding to the reflectance data. (E) SWIR reflectance curves. (F) F-value curve corresponding to the SWIR data. All data represent the mean of four biological replicates per treatment (n = 4).

### 3.9 Compound-specific hyperspectral fingerprints of *Colletotrichum* sp. P24-192

Different compounds also induced unique spectral fingerprints in *Colletotrichum* sp. P24-192, highlighting compound-specific spectral responses within the same isolate (Fig. S7). For example, in the SWIR modality, BCBGAM produced a focused diagnostic band near ≈1950 nm, whereas PhSOAM yielded a broader separation across multiple bands. These distinct spectral signatures support compound-specific spectral fingerprints under controlled in vitro conditions; no biochemical mechanism is inferred here.

## 4. Discussion

Here, we developed and evaluated a taxonomy-agnostic, multi-modal phenotyping workflow that links standardized chemical stress to reproducible morphology- and hyperspectral-based fingerprints. By integrating quantitative morphology, HSI, and supervised ML, we showed that chemical-stress phenotypes can support cross-validated prediction of crop-of-isolation labels (coffee vs cacao, in this dataset) under uniform in vitro conditions. Importantly, multi-locus phylogeny indicates mixed lineage structure among the coffee isolates, so we interpret predictive performance conservatively as evidence for a learnable phenotype signal associated with crop background within this panel, rather than proof of host adaptation or ecological memory. Together, these results position the workflow as a scalable, sensor-driven screening and decision-support approach for evaluating candidate antifungals and quantifying stress-response fingerprints.

The varied fungal responses underscored a clear structure-activity relationship dependent on both the headgroup and backbone of the four phenolic compounds. Consistent with previous reports, amide derivatives generally showed stronger antifungal activity than their fatty acid analogs. Specifically, PhSOAM, with its simple phenol and amide moiety, was the most potent growth inhibitor. This is consistent with contributions from increased polarity conferred by the amide group and steric accessibility of the unsubstituted phenol (Dembitsky, 2022; Du et al., 2015; Ōmura et al., 1974).

An unexpected observation was that BCBGFA appeared to promote growth in isolate P24 85. While this may reflect experimental variability, the consistency of this stimulation across replicates suggests a potential hormetic response (stimulation by low-dose stressors) or an isolate-specific metabolic capability to utilize the phenolic fatty acid as a carbon source (Middelhoven, 1993; Middelhoven et al., 1992). Because this study was not designed to quantify compound catabolism, we treat this observation as a hypothesis-generating result.

The divergence in morphology across isolates and compounds supports the practical idea that standardized stress assays can generate reproducible phenotype fingerprints suitable for screening and prediction. We use the term “chemical priors” as a working hypothesis to describe the possibility that prior exposure histories or persistent physiological states may shape stress-response pathways that become legible under standardized probes. In this dataset, circularity emerged as the most informative morphology-derived feature for predicting the crop-of-isolation label, whereas elongation (LWR) was not consistently expressed across the coffee panel. Notably, circularity shifts have also been observed under physical (UVC) stress in prior work (Baek et al., 2025), suggesting that certain geometric outcomes may recur across disparate stressors. However, establishing whether such recurrence reflects shared mechanisms or convergent macroscopic endpoints will require targeted molecular and experimental validation.

While linear ordination (PCA) did not separate the combined datasets, supervised ML achieved 86.7% cross-validated accuracy for classifying crop-of-isolation labels (coffee vs cacao, in this dataset), indicating that morphology contains a non-linearly separable predictive signal. Feature-importance analysis consistently identified colony circularity as the dominant predictor, with size metrics (area, perimeter) contributing additional information. When circularity was removed, classification performance remained high (86.1%), indicating that the predictive signal is distributed across multiple traits within this dataset. It is important to note that our primary cross-validation strategy involved random splitting of replicate plates, such that replicates from the same isolates could appear in both training and validation folds. Therefore, these accuracies quantify within-panel predictability rather than generalization to unseen isolates. To directly address isolate-level generalization, we additionally performed isolate-grouped cross-validation using StratifiedGroupKFold with strain as the grouping variable (*k* = 2 after excluding CGH5, due to the limited number of cacao strains; Supplementary Data 1), which provides a stricter evaluation. These results support the utility of morphology-derived features as a compact, interpretable representation for predictive screening, without requiring species-resolved inputs.

We interpret these phenotypes as plastic stress responses expressed over the 96-hour incubation window rather than the rapid selection of resistant mutants. Because the incubation timeframe and assay design do not directly test heritable change, we avoid mechanistic claims about evolutionary memory and instead report reproducible phenotype shifts under standardized conditions.

HSI complemented morphology by providing dose- and isolate-dependent spectral fingerprints under controlled imaging conditions. Prior studies indicate that VNIR/SWIR reflectance and fluorescence can differentiate physiological states in fungi, including changes associated with hydration-related bands, pigmentation, and structural components (Kong et al., 2019; Neurauter et al., 2024; Teena et al., 2014; Valenzuela-Gloria et al., 2021). In our study, the cacao-associated isolate Pestalotiopsis sp. CGH5 showed strong spectral separation across modalities, whereas the coffee isolate P24-85 exhibited a comparatively attenuated response under the same treatment series. We interpret these patterns as treatment-associated spectral variation that differentiates isolates, rather than as direct evidence for specific physiological mechanisms. Notably, the SWIR region near ∼1930 nm is moisture-sensitive, so biochemical attribution from this band alone is not warranted without explicit water-effect modeling or robustness checks.

The contrast between PhSOAM and BCBGAM in the same isolate (Fig. S7; Table S3) supports compound-specific spectral fingerprints under standardized conditions. These signatures motivate future mechanistic studies, but here we restrict interpretation to reproducible, treatment-associated spectral differences across modalities. Because the ∼1930 nm region is moisture-sensitive, future work should explicitly model or filter water effects (e.g., variable band weighting or EMSC) before attempting biochemical assignment.

These parallels between xenobiotic chemical exposure and UVC irradiation suggest that HSI can capture both compound-specific and isolate-specific physiological responses. Recurring spectral patterns across distinct stressors warrant further study, but we do not attribute them to specific ecological drivers or pathways based on the present dataset.

Crucially, the convergence of morphology and hyperspectral signatures suggests a robust, multi-modal stress-response phenotype under standardized conditions. We present isolate-dependent differences in spectral responsiveness as descriptive phenotyping outcomes that can inform screening and prioritization, while mechanistic interpretation remains future work.

The polyphyletic distribution of the coffee isolates (Fig. 3; Fig. S6) provides important context: crop-of-isolation labels are not equivalent to a single clade. While our primary headline accuracy reflects within-panel separability (replicate splitting), the supplementary grouped cross-validation (*k* = 2) and outlier-exclusion analysis (Supplementary Data 1) confirmed that the core phenotypic signal remains detectable beyond specific isolates. Thus, we interpret these results as a robust proof-of-concept for the workflow, identifying targets for future panel expansion.

Because crop-of-isolation labels are partially confounded with lineage structure in this dataset, we do not attribute the predictive phenotype signal to host adaptation as a primary driver. Instead, our results motivate future experimental designs that explicitly disentangle host background, geography, and lineage—for example, by testing multiple isolates from the same species sampled across crops and regions and evaluating isolate-blocked generalization. Within the current panel, chemical stress elicits a rich, multi-layered phenotype that is measurable by morphology and HSI and usable for predictive screening.

To synthesize these findings, Fig. 8 summarizes the end-to-end taxonomy-agnostic workflow that links standardized perturbations to reproducible fingerprints and cross-validated label prediction. In this study, the workflow links controlled chemical perturbations to reproducible morphology and spectral fingerprints and enables prediction of crop-of-isolation labels within the combined dataset. Establishing whether any component of this signal reflects heritable, environmentally induced memory (e.g., epigenetic mechanisms) will require targeted experimental designs and molecular assays beyond the scope of this work.

**Fig. 8.**
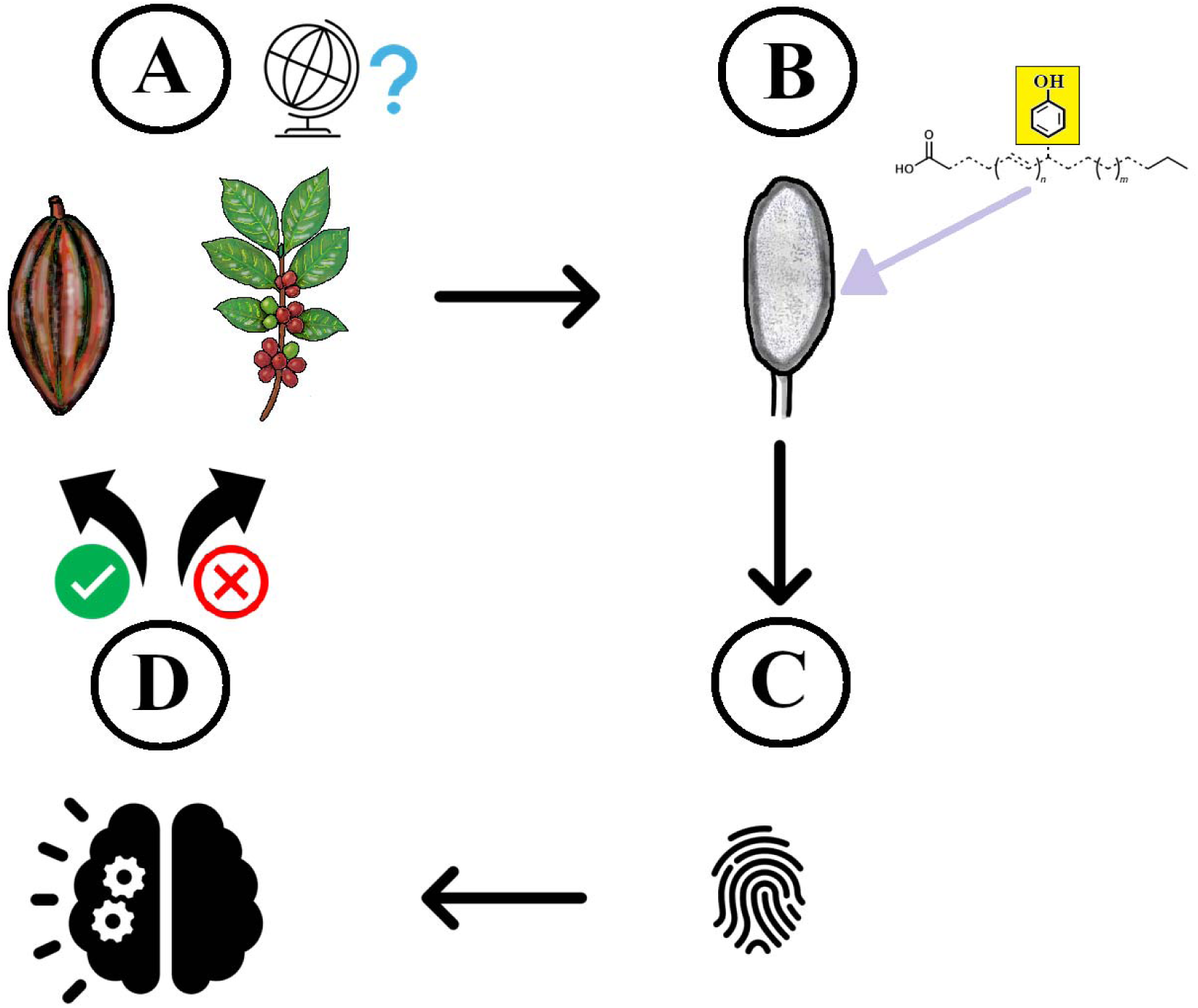
Taxonomy-agnostic phenotyping workflow for predicting crop-of-isolation labels from chemical-stress responses. (A) Sample collection and labeling. Isolates are assigned sample-source labels (e.g., crop-of-isolation) and cultured under standardized conditions. (B) Standardized perturbation and multi-modal sensing. Colonies are challenged with uniform chemical stressors and quantified using macroscopic morphology and hyperspectral imaging to generate reproducible stress-response fingerprints. (C) Feature extraction and predictiv modeling. Morphological traits and spectral summaries are used as model inputs; supervised machine learning is trained and evaluated via cross-validation to predict crop-of-isolation label within the present dataset. (D) Screening and decision support output. The workflow produce interpretable outputs (e.g., feature importance, decision thresholds) to support high-throughput screening and sensor-driven decision support. This conceptual workflow summarizes the end-to-end pipeline demonstrated here for coffee–cacao isolate panels under phenolic antifungal stress and is adaptable to other microorganisms and stressors.

Our study establishes a scalable, phenotype-driven platform for taxonomy-agnostic stress-response phenotyping and predictive modeling. The potential utility of this methodological advance is its application in high-throughput screening of eco-friendly fungicides and sensor-driven decision support for phenotype-based screening workflows. Building on this foundation, future work will prioritize: (1) validating these phenotype fingerprints in planta; (2) resolving molecular and genomic bases of stress responses; and (3) testing generality of the “chemical priors” hypothesis across diverse pathogen systems.

## 5. Conclusion

This study successfully developed and validated a high-throughput, taxonomy-agnostic phenotyping workflow that integrates quantitative morphology and hyperspectral imaging to characterize fungal stress responses. Unlike traditional multivariate statistics, which explained only ∼1.7% of the phenotypic variance, our non-linear machine learning approach achieved 86.7% classification accuracy in predicting crop-of-isolation labels within a mixed-lineage panel. Crucially, the system demonstrated significant engineering robustness; excluding the primary shape feature (circularity) resulted in a negligible performance drop (to 86.1%), confirming that the predictive fingerprint is redundantly encoded across multiple traits. This framework offers a scalable, sensor-driven solution for rapid antifungal screening and biosecurity monitoring, decoupling phenotypic profiling from the bottlenecks of strict taxonomic identification.

## CRediT authorship contribution statement

Insuck Baek: Formal analysis, Writing – review & editing. Seunghyun Lim: Formal analysis, Writing – review & editing. Amelia Lovelace: Methodology, Resources, Writing – review & editing. Sookyung Oh: Investigation, Writing – review & editing. Masoud Kazem-Rostami: Resources, Writing – review & editing. Helen Ngo: Resources, Writing – review & editing. Moon S. Kim: Funding acquisition, Writing – review & editing. Lyndel W. Meinhardt: Funding acquisition, Writing – review & editing. Lalit Kandpal: Investigation, Writing – review & editing. Minhyeok Cha: Methodology, Writing – review & editing. Chansong Hwang: Methodology, Investigation, Writing – review & editing. Richard D. Ashby: Resources, Writing – review & editing. Ezekiel Ahn: Conceptualization, Supervision, Investigation, Formal analysis, Funding acquisition, Writing – original draft, Writing – review & editing.

## Declaration of competing interest

The authors declare that they have no known competing financial interests or personal relationships that could have appeared to influence the work reported in this paper.

## Acknowledgements

Mention of any trade names or commercial products in this article is solely for the purpose of providing specific information and does not imply recommendation or endorsement by the U. S. Department of Agriculture. USDA is an equal opportunity provider and employer, and all agency services are available without discrimination. This work was supported by the U.S. Department of Agriculture, Agricultural Research Service, In-House Projects Nos. 8042-21220-258-000-D and 8072-41000-113-000-D, and by the National Institute of Food and Agriculture (NIFA) under Grant No. 2021-67021-34502. We thank Lisa Keith (Daniel K. Inouye U.S. Pacific Basin Agricultural Research Center, USDA-ARS, Hilo, Hawaii) for kindly providing the *Colletotrichum* isolates used in this study.

## Declaration of generative AI and AI-assisted technologies in the manuscript preparation process

During the preparation of this work the authors used ChatGPT (OpenAI) in order to improve language and readability. After using this tool/service, the authors reviewed and edited the content as needed and take full responsibility for the content of the published article.

## Data availability

All raw and processed data, including colony morphology measurements, per-wavelength mean spectra, and F-statistics for HSI, are provided in Supplementary Data 1. Multi-locus sequence data (ITS, GAPDH, ACT) generated for this study are openly available at figshare: https://figshare.com/articles/dataset/29924960.

## Notes

### Competing Interest Statement

The authors have declared no competing interest.

### Summary of Updates

The manuscript title has been updated to better reflect the methodological focus of the study. We have refined our terminology from 'host origin' to 'crop-of-isolation' labels to explicitly account for the polyphyletic nature of the coffee-associated isolates. Furthermore, the interpretation of 'ecological memory' has been reframed under a 'chemical priors' hypothesis, emphasizing the predictive utility of the phenotype fingerprints while remaining conservative regarding their underlying evolutionary mechanisms.

