## Supplementary Figures and Tables for "Taxonomy-agnostic hyperspectral–morphological phenotyping of fungal pathogen chemical-stress responses using machine learning"

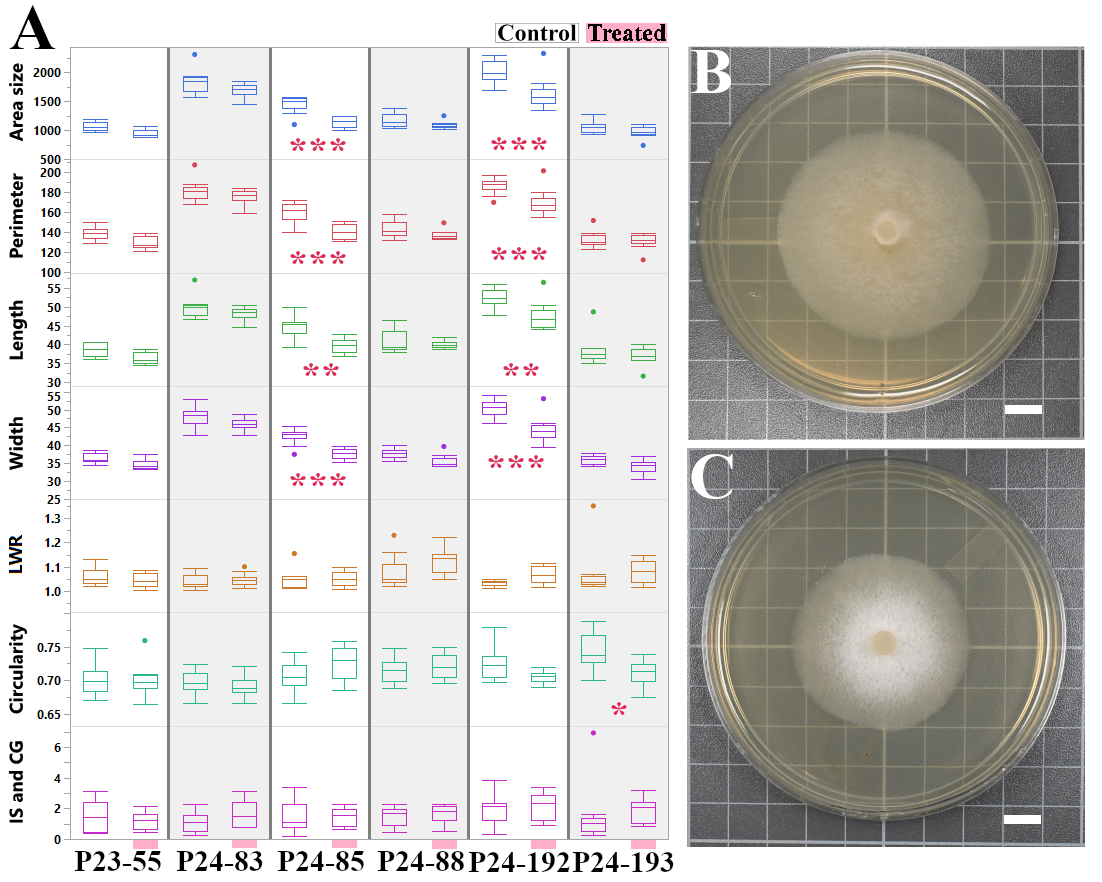

**Fig. S1. Antifungal activity and morphological disruption of six *Colletotrichum* isolates by PhSOAM.** (A) Box plots quantifying the morphological response of six *Colletotrichum* isolates (P23-55, P24-83, P24-85, P24-88, P24-192, and P24-193) to PhSOAM. For each isolate, colonies treated with PhSOAM ("Treated") are compared against untreated controls ("Control"). Measured traits include colony area (mm²), perimeter (mm), length (mm), width (mm), LWR, circularity, and an asymmetry index (IS & CG, mm). Statistically significant differences between the treated group and its corresponding control were determined by a one-way ANOVA followed by a Tukey's HSD post-hoc test. Differences are indicated by asterisks (**p* < 0.05; ***p* < 0.01; ****p* < 0.001). (B-C) Representative images of isolate P24-192 after 96 hours of growth on (B) a control (ethanol-treated) plate and (C) a plate treated with PhSOAM, illustrating the reduction in colony size. Sample size, n = 10 for all groups, except for the P24-192 and P24-193 controls (n = 11). Scale bars = 1 cm. PhSOAM exhibited the strongest overall antifungal effect among all tested compounds, with strong isolate-dependent effects on size metrics and accompanying changes in shape metrics (including LWR). However, due to the more visually striking phenotype observed with BCBGAM in P24-192 (shown in Fig. 1), PhSOAM results are presented here as a complementary Supplementary Figure.

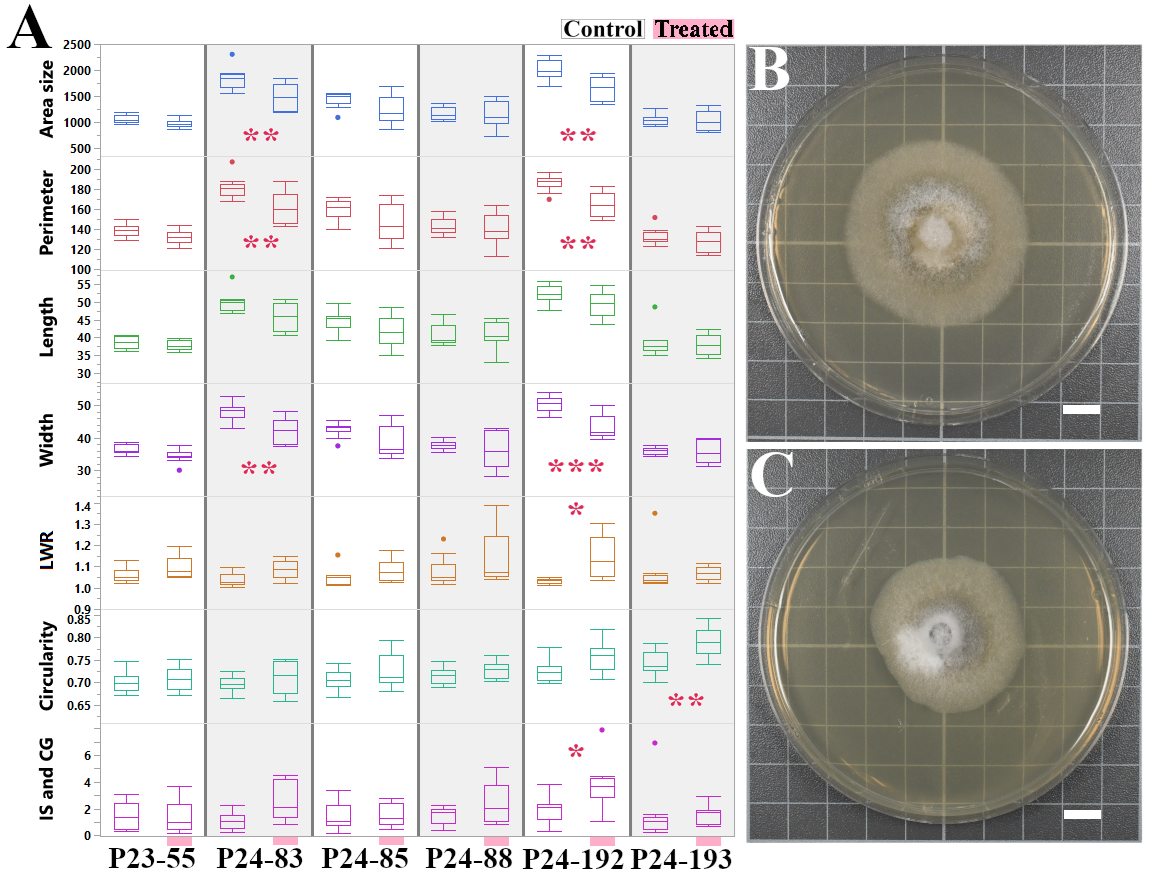

**Fig. S2. Antifungal activity of PhSOFA against coffee-associated *Colletotrichum* isolates.** (A) Box plots quantifying the morphological response of six *Colletotrichum* isolates to PhSOFA treatment compared against untreated controls. Measured traits include colony area (mm²), perimeter (mm), length (mm), width (mm), LWR, circularity, and an asymmetry index (IS & CG, mm). Significant differences between a treated group and its corresponding control (Tukey's HSD; **p* < 0.05, ***p* < 0.01, ****p* < 0.001) are marked by asterisks. (B-C) Representative images of isolate P24-83 after 96 hours on (B) a control plate and (C) a plate treated with PhSOFA, illustrating the observed reduction in colony size. Sample size, n = 10 for all groups, except for the P24-192 and P24-193 controls (n = 11). Scale bars = 1 cm.

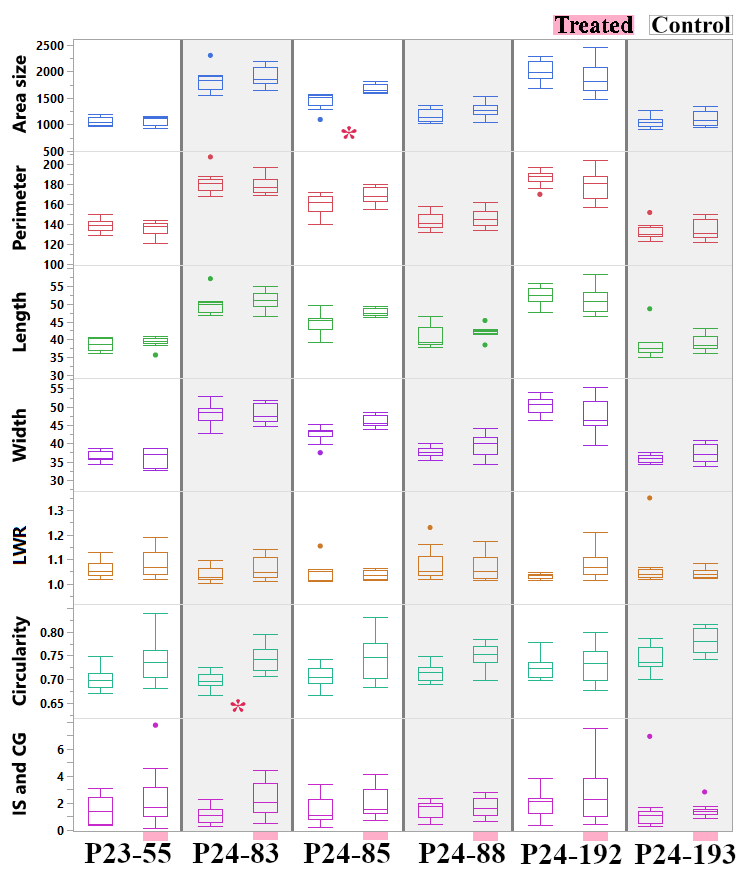

**Fig. S3. Opposing and isolate-specific effects of BCBGFA on the morphology of *Colletotrichum* isolates.** Box plots quantifying the morphological response of six *Colletotrichum* isolates to BCBGFA treatment compared against untreated controls. Measured traits include colony area (mm²), perimeter (mm), length (mm), width (mm), LWR, circularity, and an asymmetry index (IS & CG, mm). Statistically significant differences between a treated group and its corresponding control (Tukey's HSD; **p* < 0.05, ***p* < 0.01, ****p* < 0.001) are marked by asterisks. Sample size, n = 10 for all groups, except for the P24-192 and P24-193 controls (n = 11).

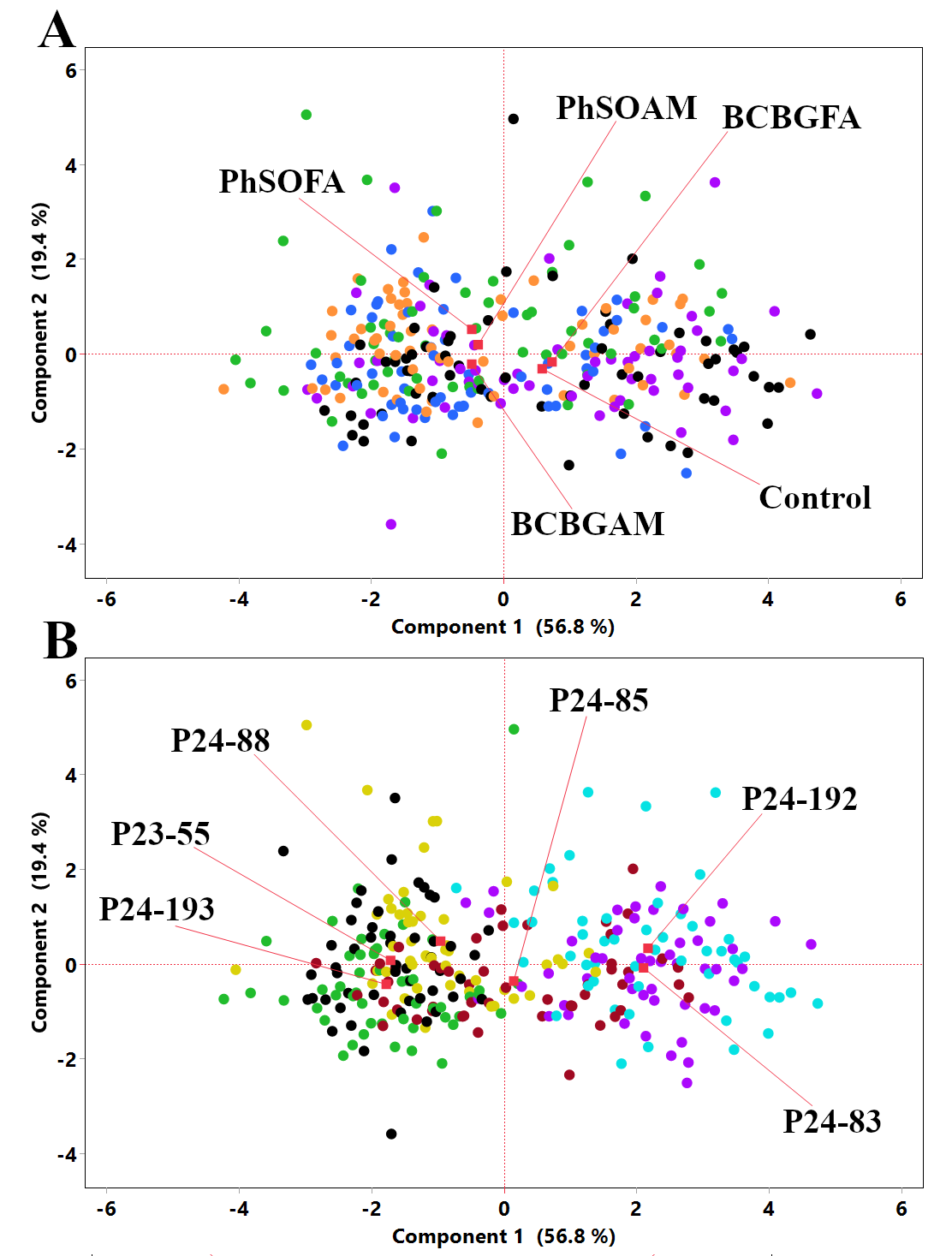

**Fig. S4. Multivariate analysis of colony morphology reveals a dominant treatment effect.** Principal Component Analysis of seven morphological traits across all four chemical treatments and six isolates. (A) PCA scores plot with samples color-coded by treatment: Control (black), PhSOAM (orange), PhSOFA (green), BCBGAM (blue), and BCBGFA (purple). (B) The identical PCA scores plot with samples color-coded by fungal isolate: P23-55 (black), P24-83 (purple), P24-85 (brown), P24-88 (yellow), P24-192 (cyan), and P24-193 (green). The red squares indicate the mean of each treatment group in (A) and each fungal isolate group in (B). PERMANOVA detected a small but significant treatment effect (pseudo-F = 4.06, R² = 0.0518, *p* = 0.001) and a stronger isolate effect (pseudo-F = 13.00, R² = 0.180, *p* < 0.0001), yet both factors showed substantial group overlap in PCA space.

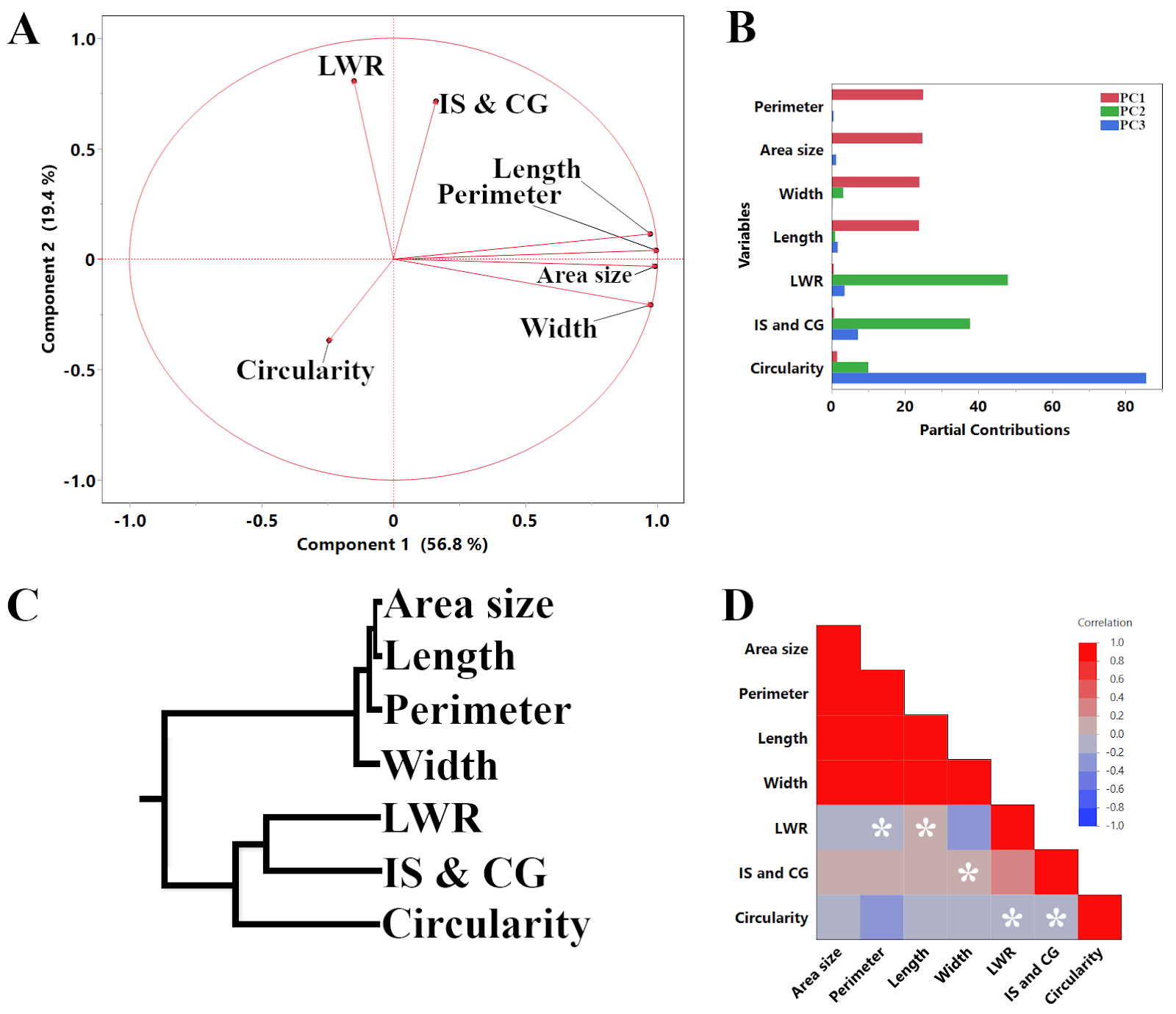

**Fig. S5. Analysis of morphological trait relationships.** (A) PCA loading plot illustrating the contribution of each of the seven log-transformed morphological traits to the first two principal components. (B) Partial contributions plot showing the percentage that each principal component contributes to the variance of each trait. (C) Hierarchical clustering dendrogram of the characteristics using Ward's method, showing a clear grouping of size-related versus shape-related metrics. (D) Pearson's correlation matrix of all traits. Red indicates a positive correlation, blue indicates a negative correlation, and white asterisks (*) denote correlations that are not statistically significant (*p* > 0.05).

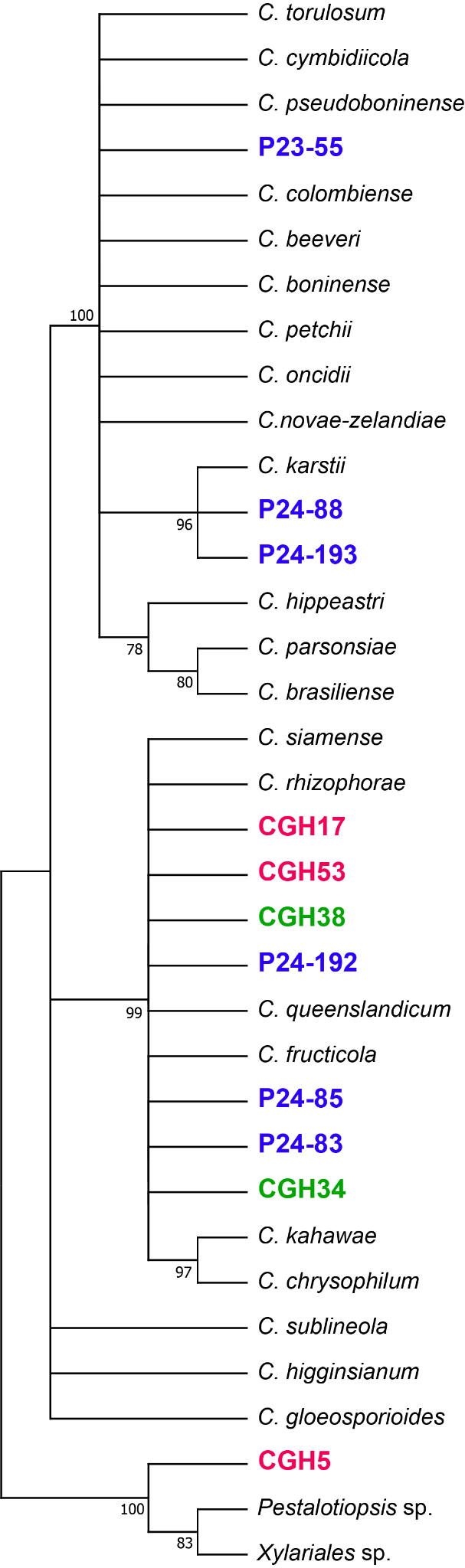

**Fig. S6. ITS phylogeny reveals close genetic relationships between isolates from coffee and cacao crop-associated panels.** The phylogenetic tree was inferred by the Neighbor-Joining method using the ITS region. The evolutionary distances were computed using the Kimura 2-parameter (K2+G) model. Numbers at the nodes represent bootstrap support values based on 1,000 replications (≥75% shown). Isolates from coffee are highlighted in blue, isolates from cacao in red, and isolates from associated shade trees in green. The tree demonstrates that some coffee-associated isolates (e.g., P24-192, P24-85, P24-83) are phylogenetically intermixed with isolates from cacao (CGH17, CGH53) and shade trees (CGH34, CGH38) within a single, strongly supported clade (99% bootstrap).

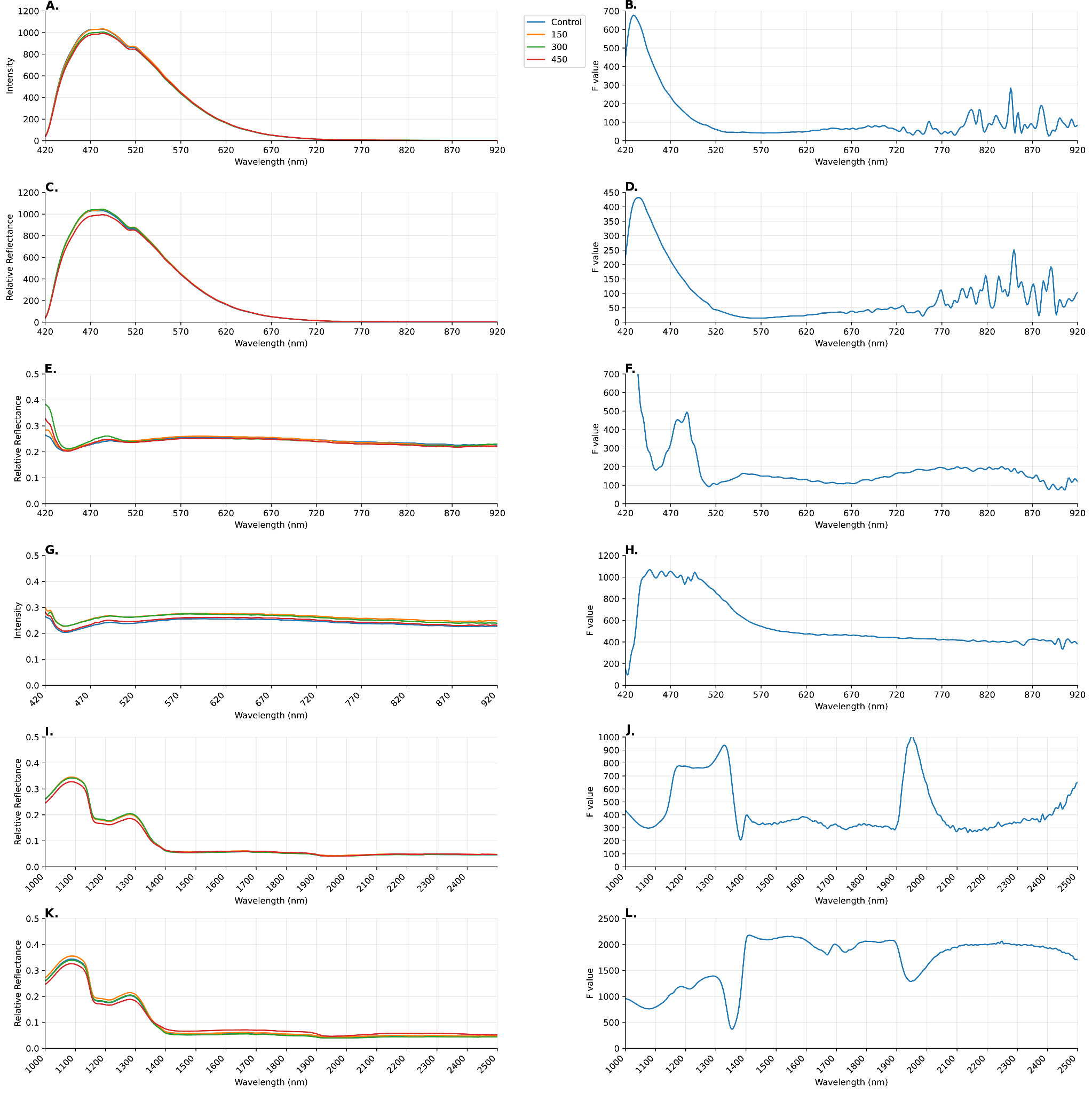

**Fig. S7. Compound-specific hyperspectral signatures of *Colletotrichum* sp. P24-192 treated with BCBGAM and PhSOAM across three spectral modalities.** Panels show colonies treated with 0 µg (control, blue), 150 µg (orange), 300 µg (green), and 450 µg (red) of each compound (n = 4 per group). (A) Fluorescence intensity profiles for BCBGAM treatment. (B) F-value curve for BCBGAM fluorescence. (C) Fluorescence intensity profiles for PhSOAM treatment. (D) F-value curve for PhSOAM fluorescence. (E) Reflectance intensity curves for BCBGAM treatment. (F) F-value curve for BCBGAM reflectance. (G) Reflectance intensity curves for PhSOAM treatment. (H) F-value curve for PhSOAM reflectance. (I) SWIR reflectance curves for BCBGAM treatment. (J) F-value curve for BCBGAM SWIR. (K) SWIR reflectance curves for PhSOAM treatment. (L) F-value curve for PhSOAM SWIR. All data represent the mean of four biological replicates per treatment (n = 4).

**Table S1. Performance of machine learning models for classifying crop-of-isolation labels.**

| Method | TP | FN | FP | TN | Sensitivity | Specificity | Precision | Accuracy | F1 | MCC |
| --- | --- | --- | --- | --- | --- | --- | --- | --- | --- | --- |
| Decision Tree | 139 | 41 | 22 | 158 | 0.772 | 0.878 | 0.863 | 0.825 | 0.815 | 0.654 |
| Bootstrap Forest | 147 | 33 | 25 | 155 | 0.817 | 0.861 | 0.855 | 0.839 | 0.835 | 0.678 |
| Boosted Tree | 145 | 35 | 28 | 152 | 0.806 | 0.844 | 0.838 | 0.825 | 0.822 | 0.651 |
| Naive Bayes | 144 | 36 | 66 | 114 | 0.800 | 0.633 | 0.686 | 0.717 | 0.739 | 0.440 |
| Neural Boosted | 155 | 25 | 23 | 157 | 0.861 | 0.872 | 0.871 | 0.867 | 0.866 | 0.733 |
| Support Vector Machines | 148 | 32 | 26 | 154 | 0.822 | 0.856 | 0.851 | 0.839 | 0.836 | 0.678 |
| Fit Stepwise | 150 | 30 | 28 | 152 | 0.833 | 0.844 | 0.843 | 0.839 | 0.838 | 0.678 |
| Nominal Logistic | 150 | 30 | 25 | 155 | 0.833 | 0.861 | 0.857 | 0.847 | 0.845 | 0.695 |
| Generalized Regression Lasso | 148 | 32 | 21 | 159 | 0.822 | 0.883 | 0.876 | 0.853 | 0.848 | 0.707 |

The table summarizes key performance metrics derived from the 5-fold cross-validation for the nine algorithms trained on the balanced, log-transformed morphological dataset. Metrics shown include the number of true positives (TP), false negatives (FN), false positives (FP), and true negatives (TN), as well as calculated values for Sensitivity, Specificity, Precision, Accuracy, F1, and Matthews Correlation Coefficient (MCC).

**Table S2. Relative importance of predictor variables for crop-of-isolation label classification.**

| **Trait** | **Main Effect** | **Total Effect** |
| --- | --- | --- |
| Circularity | 0.383 | 0.592 |
| Area size | 0.223 | 0.405 |
| Perimeter | 0.063 | 0.168 |
| Treatment (Phenol) | 0.036 | 0.119 |
| Width | 0.043 | 0.11 |
| Length | 0.046 | 0.109 |
| LWR | 0.015 | 0.043 |
| IS & CG | 0.01 | 0.022 |

The table shows the feature importance metrics derived from the Neural Boosted model on the log-transformed dataset. Predictors are ranked by their "Importance," which indicates their overall contribution to the model's ability to distinguish between the cacao and coffee crop-of-isolation labels in the combined dataset.

**Table S3. Summary of hyperspectral responses observed in selected fungal pathogens following treatment with phenolic-branched fungicides.**

| Isolate | Compound | Modality | Wavelength(s) (nm) | Spectral Response Description |
| --- | --- | --- | --- | --- |
| CGH5 | PhSOAM | Fluorescence | ≈470–560 (peak ~510 nm) | Dose-dependent separation across doses. |
| CGH5 | PhSOAM | VNIR (Reflect.) | ≈430–560 | Treatment-associated spectral variation across doses. |
| CGH5 | PhSOAM | SWIR | ≈1460–1470; ≈1900–2000 | Treatment-associated spectral variation in moisture-sensitive SWIR regions; interpretation limited without explicit water-effect modeling. |
| P24-85 | PhSOAM | Fluorescence | ≈450–560 (peak ~470–510) | Non-monotonic dose pattern. |
| P24-85 | PhSOAM | VNIR (Reflect.) | ≈430–490 (prominent near ≈485 in mean spectra) | Dose-specific variation, most pronounced at 150 µg. |
| P24-85 | PhSOAM | SWIR | ≈1400–1500; ≈1900–2050 | Attenuated/flat spectral response across doses relative to CGH5 in moisture-sensitive SWIR regions. |
| P24-192 | PhSOAM | SWIR | ≈1400–1900; ≈2250; ≈1290 | Broad treatment-associated spectral variation across SWIR; secondary features near ≈2250 and ≈1290. |
| P24-192 | BCBGAM | SWIR | ≈1950 (± ~15); ≈1180–1350 (max ≈1326) | Focused treatment-associated spectral variation near ≈1950 with additional variation across ≈1180–1350 (descriptive; directionality not assigned). |

Reported spectral regions include moisture-sensitive SWIR bands (≈1900–2050 nm); interpretations are limited to treatment-associated spectral variation without biochemical assignment.

**Table S4. Information for coffee-associated *Colletotrichum* spp. isolates used in this study.**

| Isolate ID | Crop-of-isolation label (*C. arabica*) | Host Tissue | Location (Hawaii) | Year of Isolation |
| --- | --- | --- | --- | --- |
| P23-55 | cv. Obata | Leaf spots | Holualoa | 2023 |
| P24-83 | cv. Typica | Leaf spots | Hilo | 2024 |
| P24-85 | cv. Typica | Leaf spots | Hilo | 2024 |
| P24-88 | cv. Catimor | Necrotic stems | Hāna | 2024 |
| P24-192 | cv. Bourbon | Necrotic stems | Hilo | 2024 |
| P24-193 | cv. Bourbon | Necrotic stems | Hilo | 2024 |

All coffee-associated isolates were provided by the U.S. Department of Agriculture, Agricultural Research Service collection in Beltsville, MD. The isolates were originally collected between 2023 and 2024 from various cultivars and tissues of *Coffea arabica* at different locations in Hawaii, USA.

Note: Full statistical results, including ANOVA *p*-values for all trait comparisons and detailed machine learning robustness checks (e.g., feature ablation results), are provided in the accompanying Excel file (Supplementary Data 1).
